# A Thin-Film Lubrication Model for Biofilm Expansion Under Strong Adhesion

**DOI:** 10.1101/2021.11.16.468738

**Authors:** Alexander K. Y. Tam, Brendan Harding, J. Edward F. Green, Sanjeeva Balasuriya, Benjamin J. Binder

## Abstract

Understanding microbial biofilm growth is important to public health, because biofilms are a leading cause of persistent clinical infections. In this paper, we develop a thin-film model for microbial biofilm growth on a solid substratum to which it adheres strongly. We model biofilms as two-phase viscous fluid mixtures of living cells and extracellular fluid. The model tracks the movement, depletion, and uptake of nutrients explicitly, and incorporates cell proliferation via a nutrient-dependent source term. Notably, our thin-film reduction is two-dimensional and includes the vertical dependence of cell volume fraction. Numerical solutions show that this vertical dependence is weak for biologically-feasible parameters, reinforcing results from previous models in which this dependence was neglected. We exploit this weak dependence by writing and solving a simplified one-dimensional model that is computationally more efficient than the full model. We use both the one and two-dimensional models to predict how model parameters affect expansion speed and biofilm thickness. This analysis reveals that expansion speed depends on cell proliferation, nutrient availability, cell–cell adhesion on the upper surface, and slip on the biofilm–substratum interface. Our numerical solutions provide a means to qualitatively distinguish between the extensional flow and lubrication regimes, and quantitative predictions that can be tested in future experiments.

## 1 Introduction

Biofilms are sticky communities of micro-organisms that exist on surfaces. An estimated 80% of all microbes on Earth exist in biofilm colonies [1], and they have extensive effects on human life. Most notably, biofilms are responsible for up to 80% of bacterial infections [2]. Since biofilms are highly resistant to anti-microbial therapy [3–5], treating these infections often requires expensive and dangerous surgery [1], generating a large health burden in public hospitals. Another negative consequence of biofilms is dental plaque [6], but biofilms can also be beneficial when harnessed for wastewater treatment [7] and fuel production [8]. In all of these scenarios, it would be advantageous to predict and control biofilm growth, for example to minimise the spread of an infection. In addition to their effects on human life, biofilms have also fascinated scientists for their ability to form diverse and complex patterns [9]. Owing to their importance to infection control and the understanding of collective cell behaviour, biofilms remain widely studied today.

We develop and solve a thin-film lubrication model for biofilm formation. Our model is based on mat formation experiments of the budding yeast *Saccharomyces cerevisiae* [10, 11], which consist of cells spreading radially on an agar substratum from which they obtain nutrients. Previously, we developed an extensional flow model to investigate growth by sliding motility [12]. However, biofilm formation in practice also depends on the ability of cells to adhere to surfaces [13]. In particular, the invasiveness of an infection depends on the biofilm’s ability to adhere to living tissue or indwelling devices such as catheters, stents, and prostheses [2, 14, 15], and subsequently proliferate. Models for strongly-adhesive biofilms commonly involve the lubrication scaling regime and a no-slip condition on the biofilm–substratum interface. In this work, we apply the lubrication scaling to the general model introduced in Tam et al. [12].

The paper is structured as follows. In §1.1, we review relevant biological literature and previous models with strong biofilm–substratum adhesion. Based on this, we adapt our previous two-phase model to a relevant lubrication regime of comparatively high pressure and surface tension in §2, and present the thin-film equations. Unlike many previous lubrication models, our governing equations are two-dimensional. However, numerical solutions to a regularised system reveals that vertical dependence on cell volume fraction is weak for biologically-relevant parameters. In §3, we exploit this by deriving and solving a computationally-efficient one-dimensional version of our model, in which we neglect this vertical dependence. We then use both the full two-dimensional and simplified one-dimensional models to investigate the effect of parameters on expansion in §4. This analysis quantifies how a balance between cell proliferation, surface tension forces, and biofilm–substratum slip governs the size and shape of a biofilm. We close the paper with a discussion and conclusion in §5.

### 1.1 Biological Background and Previous Models

A wide range of mathematical models exist for biofilms [16]. These include discrete or hybrid models, which track the movement of individual cells [17, 18]. Another approach is to treat cells and nutrients as continuous density fields, and model their movement and interactions with reaction–diffusion equations [9, 11]. In this paper, we focus instead on continuum mechanical models. Cells in biofilm colonies interact mechanically with each other, and a self-produced extracellular matrix (ECM) in which cells reside. This ECM consists of water and extracellular polymeric substances (EPS), and helps the biofilm survive by facilitating nutrient transport and providing a physical barrier to harmful substances [19, 20]. Continuum mechanical models provide a way to investigate how the combination of nutrient-dependent cell proliferation, and mechanical interactions between cells and their environment, affect biofilm growth.

A class of fluid mechanical models was introduced by Wanner and Gujer [21]. These models involve biofilms growing vertically on solid, non-reactive substrata, and incorporate hydrodynamics of a nutrient-rich liquid culture medium in which the biofilm is immersed [22]. However, since these models treated the biofilm as rigid, they did not incorporate the mechanics of the cells and ECM. Furthermore, in this work we consider biofilms spreading radially over a substratum from which they obtain nutrients. This is relevant to biofilm formation experiments, as well as biofilm growth on living tissue. For these reasons, we do not consider models of vertical growth in detail, and refer readers to the review by Klapper and Dockery [23] for further information.

For spreading biofilms, a common approach is to model biofilm constituents themselves as fluids. The differential adhesion hypothesis (DAH) of Steinberg [24] introduced the idea of treating collections of cells as a viscous fluid. According to the DAH, cell populations behave as viscous liquids, whereby adhesive and cohesive interactions between cells are analogous to surface tension [25]. Similarly, some authors drew parallels between diffusion-limited aggregation and Hele-Shaw flow [26, 27], hypothesising that colonies can be modelled as viscous fluids. These observations have since been validated in experiments showing that biofilms behave as viscous fluids on time scales longer than the order of seconds [28, 29], with Reynolds numbers of Re < 1 × 10^−3^ [30]. Furthermore, experiments by Epstein et al. [20] showed that spreading *Bacillus subtilis* biofilms are highly resistant to liquid and gas penetration. The authors speculated that biofilm surface tension assists this de-wetting phenomenon. Preceding work by Angelini et al. [31] also suggested that biofilm expansion depends on both surface tension and the surface tension gradient. This motivates lubrication regime models, in which pressure can vary across the biofilm–air interface.

Multi-phase models provide a way to distinguish between active cells and passive fluid, such that active and passive matter are modelled as separate viscous fluids with their own behaviour and properties. A detailed framework for the construction of multi-phase models was provided by King and Oliver [32]. In these models, biofilms are typically modelled as multi-phase mixtures of cells, EPS, and external liquid [33–40]. Applying conservation of mass and momentum for each fluid phase then enables the mechanical behaviour of each fluid, and interactions between phases, to be taken into account.

Osmotic swelling can be an important mechanism in biofilm expansion [41–43]. According to Yan et al. [42], extracellular matrix production gives rise to an osmotic pressure gradient across the interface between the biofilm and external environment. Bacterial biofilms, for example those of *B. subtilis*, commonly exhibit ECM fractions of 50–90% [16], and this can be as high as 95–98% [40, 44]. Since these large quantities of ECM give rise to larger osmotic pressure gradients, osmotic swelling is likely to be an important mechanism. However, in some biofilms this ECM fraction is much smaller. For example, we observed the ECM fraction to be approximately 10% in biofilms of the yeast *S. cerevisiae* [12]. In these colonies, osmotic swelling is likely to be less important compared to cell proliferation in facilitating expansion. Therefore, although this assumption will not apply to all biofilms, we neglected osmotic swelling in this work. This enables comparison with the extensional flow results reported in Tam et al. [12], in which we neglected osmotic swelling to focus on expansion driven by sliding motility.

Once the model has been established, the resulting system of equations is often complicated. A common technique is to simplify the model by exploiting the thin geometry of a spreading biofilm. In many previous works that adopted this thin-film approximation in multi-phase models, the authors derived a fourth-order lubrication equation in one spatial dimension for the biofilm height [31, 41, 43, 45–49]. However, applying the lubrication thin-film reduction to the model of Tam et al. [12] yields a two-dimensional system of equations, unless we assume that cell volume fraction is independent of *z* from the outset. As per the model of Ward and King [46], this assumption of *z*-independence is justified for early biofilm growth, during which cell volume fraction can be assumed constant because ECM production is negligible. In contrast, we aim to model the transition from early growth to biofilm maturity, during which the volume fraction of extracellular fluid varies with space and time. Our first objective is thus to apply the lubrication scaling regime to our previous work [12], and investigate whether the commonly-made assumption that volume fraction is independent of depth [32, 37, 41, 45–48] is still valid beyond early growth. We will then use our model to obtain a quantitative understanding of how biofilm growth depends on the balance between nutrient depletion and uptake, cell proliferation, and mechanical forces.

## 2 Axisymmetric Thin-Film Lubrication Model

We consider three-dimensional growth of biofilms that adhere strongly to a solid substratum. Like Tam et al. [12], we assume radial symmetry from the outset, and illustrate this in Figure 2.1. The biofilm has the characteristic height *H*_*b*_ and the characteristic radius *R*_*b*_, and is bounded below by a solid substratum of depth *H*_*s*_, and above by a free surface *h*(*r, t*). We refer to the leading edge of the biofilm, *S*(*t*), as the contact line. To model nutrient uptake, we introduce the nutrient concentration in the substratum *g*_*s*_(*r, z, t*), defined for −*H*_*s*_ < *z* < 0. As the biofilm grows, nutrients can enter the biofilm across the *z* = 0 interface, and we denote the concentration of nutrients in the biofilm as *g*_*b*_(*r, z, t*), defined for 0 < *z* < *h*(*r, t*) and 0 < *r* < *S*(*t*).

**Figure 2.1:**
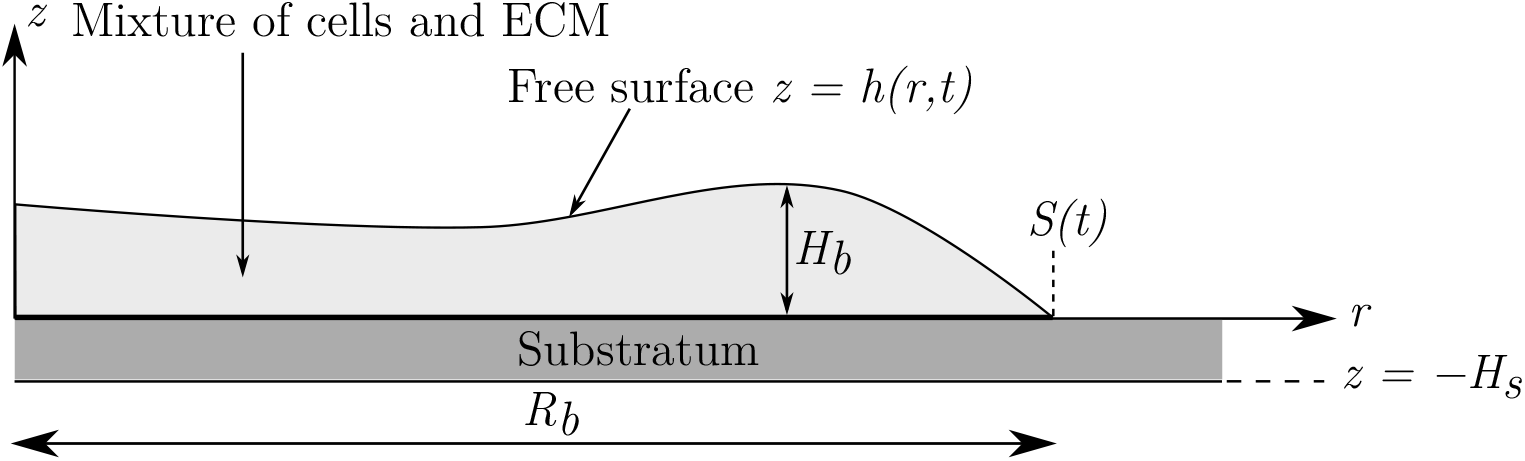
Simplified representation of a vertical slice through the centre of the biofilm and substratum [12]. The biofilm exists in the region 0 < *z* < *h*(*r, t*), where 0 < *r* < *S*(*t*).

We adopt a two-phase description of the biofilm. These phases consist of a living cell phase that actively contributes to biomass production, and a phase consisting of the ECM and other passive constituents. We define the (spatially-averaged) volume fractions of the living cells and ECM phases to be *ϕ* _*n*_(*r, z, t*) and *ϕ*_*m*_(*r, z, t*), respectively, and assume no-voids, that is *ϕ*_*n*_ + *ϕ*_*m*_ = 1. The multi-phase nature of the model enables us to track the distribution of living cells in the biofilm, which determines the location of subsequent cell proliferation, ECM production, and cell death. To simplify the model, we assume large drag between fluid phases [50], such that both fluid phases move with the common velocity ***u*** = (*u*_*r*_, *u*_*z*_). Since cells and the extracellular matrix are both primarily composed of water, we also assume the phases have the same pressure, *p*, and dynamic viscosity, *μ* [50]. The Newtonian viscous stress tensor, ***σ***, then describes the mechanics of the bulk mixture. This simplifies the model further, eliminating the need for separate momentum balance equations for each phase.

Given these assumptions, the general mass and momentum balance equations for our model are

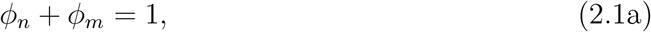

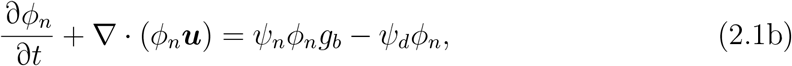

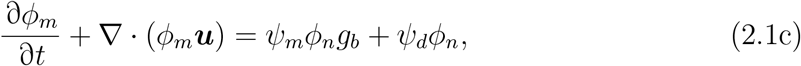

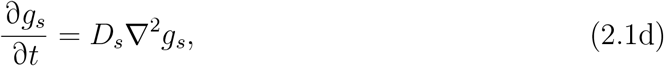

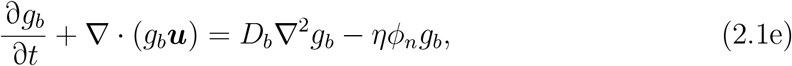

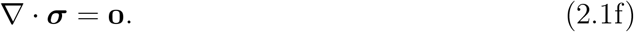

These equations differ from Tam et al. [12] only in that we allow nutrients in the biofilm to advect with both fluid phases, as opposed to assuming that the nutrients advect with the extracellular fluid phase only. In the mass balance equations, we adopt linear growth kinetics, which are the simplest forms that model cell proliferation proportional to cell volume fraction and nutrient concentration, and cell death proportional to local volume fraction. We also assume that dead cells immediately become part of the extracellular phase. The constants *ψ*_*n*_, *ψ*_*m*_, and *ψ*_*d*_ in the source terms are the cell production rate, ECM production rate, and cell death rate, respectively. In the nutrient mass balance equations, the parameters *D*_*s*_ and *D*_*b*_ are the diffusivities of nutrients in the substratum and biofilm, respectively, and *η* is the nutrient consumption rate. We assume that nutrients disperse by diffusion in the substratum, and by both diffusion and advection inside the biofilm, where they can also be consumed by cells. To obtain the momentum balance equation (2.1f), we adopt a Stokes flow description and neglect inertial and body forces. In axisymmetric cylindrical geometry, the relevant components of the stress tensor, ***σ***, are

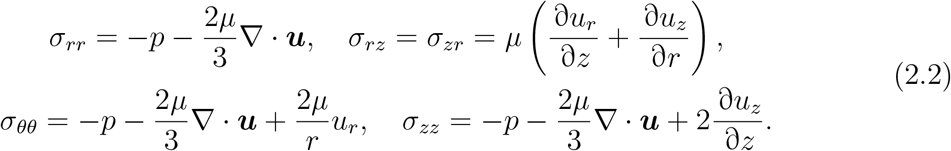

Due to cell proliferation and death, the stress components (2.2) retain divergence terms that would otherwise vanish due to incompressibility. Invoking Stokes’ hypothesis, we adopt the standard coefficient −2*μ/*3 for these divergence terms [39, 46]. We also neglect growth pressure due to cell–cell contact, which was previously considered in similar models [51, 52]. Instead, we assume that microbes cannot respond actively to chemical or mechanical cues from the environment. This is appropriate for yeasts because they are non-motile, and also bacteria because they often lose swimming motility in biofilm environments [39]. Instead of this growth pressure, we suggest that material incompressibility is sufficient to drive expansion when cells proliferate.

To obtain general boundary conditions, we first assume that nutrients cannot pass through the base of the substratum or the biofilm–air interface. Nutrients can cross the biofilm–substratum interface, with a flux proportional to the local concentration difference across the interface. For the fluids, we assume that both phases exist in the biofilm only and cannot pass through the biofilm–substratum interface, on which we also impose a general slip condition. Finally, we impose the usual kinematic and zero tangential stress conditions on the free surface, and assume that free surface normal stress is proportional to mean local curvature. The complete boundary conditions are then

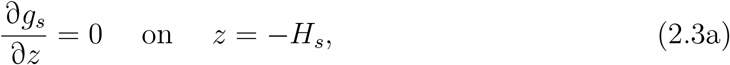

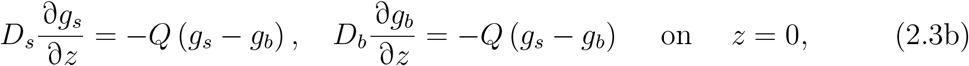

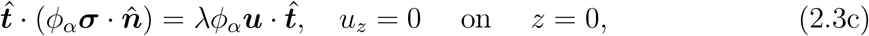

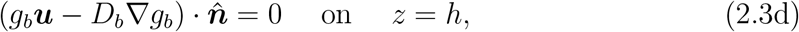

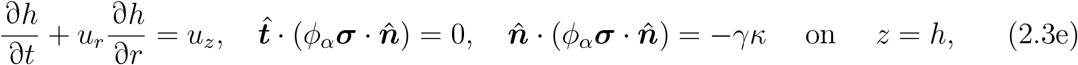

where *Q* is the mass transfer coefficient for nutrients, *λ* is a coefficient representing the strength of fluid–substratum adhesion (assumed to be the same for both phases), 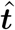 and 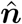 are unit tangent and normal vectors respectively, *γ* is the surface tension coefficient, and 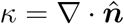 is the mean free surface curvature. In biological contexts, this surface tension represents the strength of cell–cell adhesion on the free surface [25].

Unlike extensional flows, with strong adhesion pressure can vary across the biofilm–air interface, such that the normal force due to the pressure difference balances with surface tension. To capture this, we adopt a different pressure scaling to Tam et al. [12], such that the pressure balances with surface tension terms. Defining the biofilm aspect ratio to be *ε* = *H*_*s*_*/R*_*b*_ ≪ 1 such that *H*_*b*_ ∼ 𝒪 (*ε*), we introduce the scaled variables

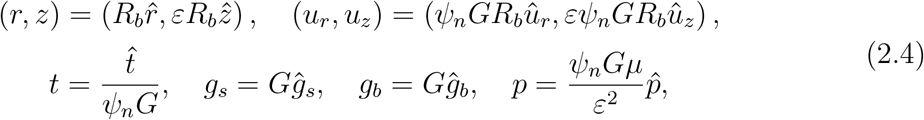

where *G* is the initial nutrient concentration in the substratum. We then exploit the thin biofilm aspect ratio to obtain a simplified leading-order system of governing equations. Expanding variables in powers of *ε*^2^ and applying the thin-film reduction, we obtain the leading-order governing equations (where variables are now expressed as dimensionless leading-order quantities)

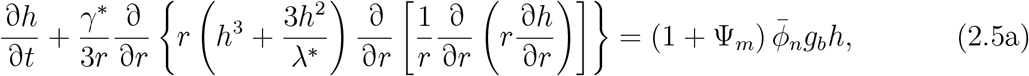

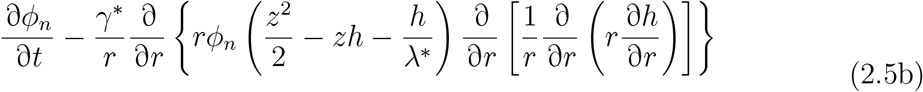

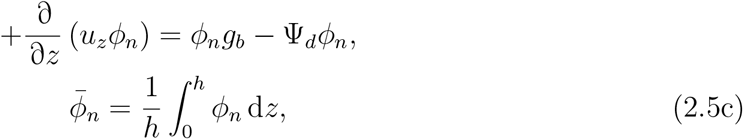

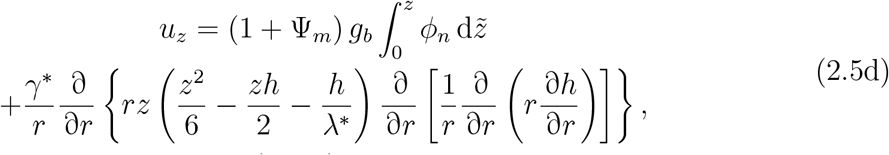

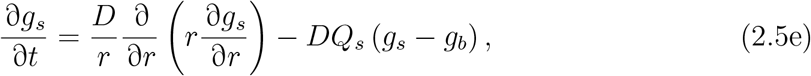

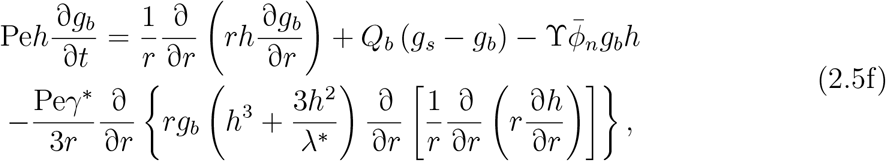

where the dimensionless parameters are

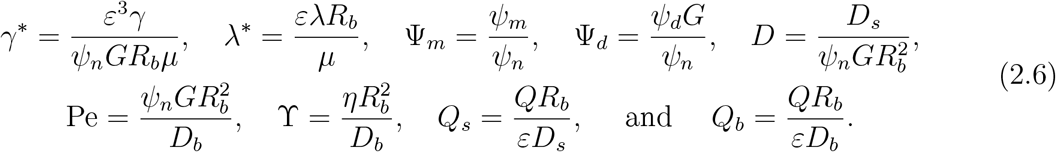

To obtain the system (2.5), we use the no-voids condition (2.1a) to eliminate *ϕ*_*m*_, such that (2.5a) is the mass conservation equation for *ϕ*_*n*_ + *ϕ*_*m*_. We subsequently obtain explicit formulae for the pressure and leading-order radial velocity *u*_*r*_, and substitute these into the mass balance equations. Full details on the thin-film reduction used to derive (2.5) and (2.6) are provided in the Supplemental Material. The system (2.5) constitutes the axisymmetric thin-film lubrication model for the biofilm height *h*(*r, t*), cell volume fraction *ϕ*_*n*_(*r, z, t*), fluid velocity *u*_*z*_(*r, z, t*), and nutrient concentrations *g*_*s*_(*r, t*) and *g*_*b*_(*r, t*). Importantly, in our thin-film reduction it is not possible to eliminate the *z*-dependence in cell volume fraction, because cells advect with the fluid velocity *u*_*r*_, which is *z*-dependent. This distinguishes our model from previous approaches that either treat the cell fraction as constant [32, 37, 46], or independent of *z* from the outset [41, 43, 45, 47, 48].

### 2.1 Precursor Film Regularisation

Imposing a no-slip condition on the biofilm–substratum interface prevents biofilm expansion in the absence of suitable regularisation. This apparent paradox is commonly encountered in models involving a lubrication equation [41, 46, 53, 54]. One method for dealing with moving contact lines is to introduce a precursor film [54]. This is an artificial thin layer of fluid with thickness *b* ≪ 1 existing ahead of the biofilm front. A physical interpretation of this is to represent the characteristic scale of surface roughness in the agar [53]. Following Ward and King [46], we adopt this precursor film to regularise the model. We assume that the precursor layer consists entirely of passive fluid (no cells), and that nutrient uptake or consumption does not occur in the precursor film. Mathematically, this involves modifying the model equations to extinguish the relevant terms wherever *h* ≤ *h*\*, for some *h*\* ≥ *b* [54]. The regularised model is then

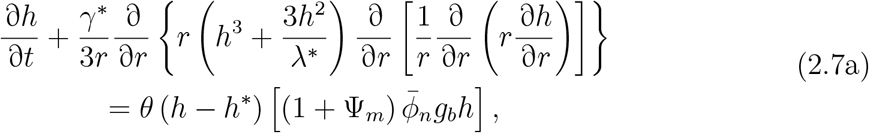

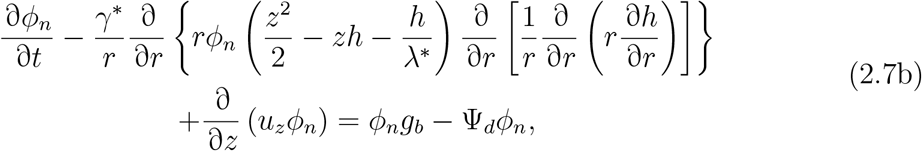

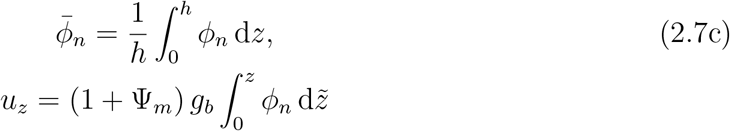

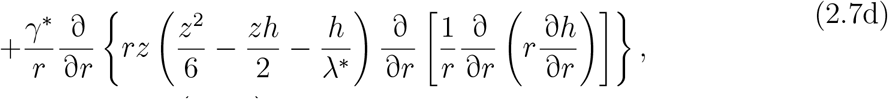

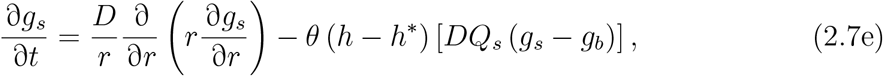

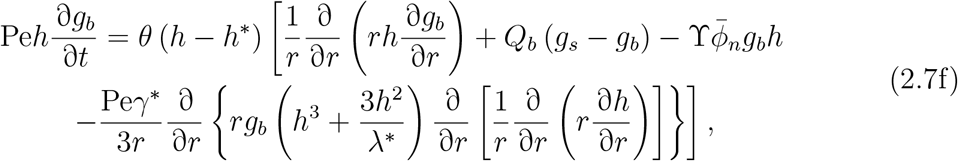

where *θ* denotes the Heaviside step function. The formulation (2.7) assumes that cells are not present without sufficient biofilm thickness to support them, and that the biofilm cannot take up nutrients in regions with no cells. The constant *h*\* represents the dimensionless thickness of a single cell. Given that the diameter of yeast cells is approximately 4 μm, and the characteristic biofilm height is approximately 2 mm, the value *h*\* = 0.002 is appropriate. Computing numerical solutions of the no-slip lubrication model then involves solving the regularised system (2.7), subject to appropriate initial and boundary conditions.

### 2.2 Initial and Boundary Conditions

We obtain initial conditions for the regularised thin-film equations (2.7) by assuming that the substratum is initially filled uniformly with nutrients, and that no nutrients are yet present in the biofilm. For *h*(*r*, 0), we require a function such that there is a defined region with *h* = *b* ahead of the biofilm, and that relevant higher derivatives of *h* with respect to *r* are continuous throughout the entire domain. For this purpose, we modify the parabolic initial condition of Tam et al. [12] in the same way as Ward and King [46]. Finally, for the cell volume fraction *ϕ*_*n*_(*r, z*, 0), we choose a polynomial form such that *ϕ*_*n*_ = 0 in the precursor film, and that the first derivatives of *ϕ*_*n*_ with respect to *r* and *z* are continuous throughout the domain. Although this is not necessarily physical, it provides an initial volume fraction profile that varies with *r* and *z*. The initial conditions are then

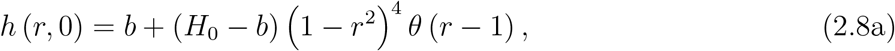

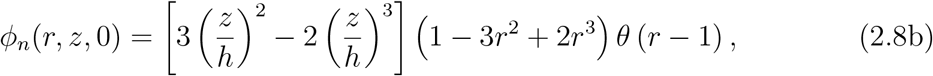

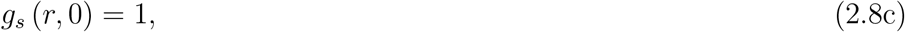

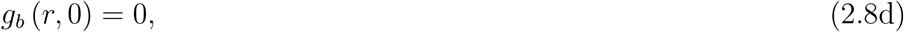

where *H*_0_ is the initial height of the biofilm at its centre. The conditions (2.8) set the initial contact line position to *S*(0) = 1, and satisfy *h* = *b* for *r* ≥ 1 and *ϕ*_*n*_ = 0 for *r* ≥ 1.

For the boundary conditions, we assume radial symmetry at *r* = 0 in all variables. Therefore, we obtain the conditions

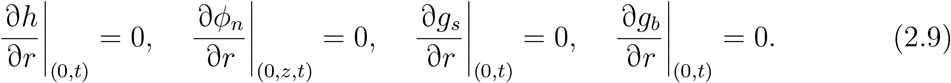

We also assume that the centre of the biofilm is fixed, and that fluid cannot cross *r* = 0. Thus, the radial component of the fluid velocity must then be zero there, that is *u*_*r*_ (0, *z, t*) = 0. The velocity *u*_*r*_ is known explicitly, and on applying L’Hôpital’s rule we obtain

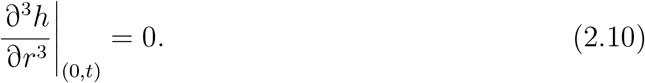

At the edge of the Petri dish, we impose no-flux conditions for the nutrient concentrations in both the substratum and biofilm. Owing to the precursor regularisation, we also have that the biofilm height is fixed at *h* = *b*, and that the precursor film has a constant height over its extent. This yields the conditions

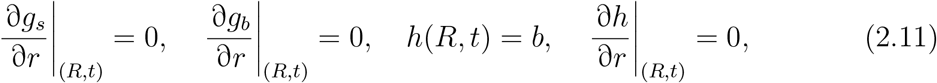

where *r* = *R* is the right-hand boundary of the domain. We now have a closed, axisymmetric model consisting of the system (2.7), initial conditions (2.8), and boundary conditions (2.9)–(2.11).

### 2.3 Parameters

In Table 2.1, we present default model parameters to be used throughout. In all numerical solutions, we choose a dimensional cell production rate of *ψ*_*n*_ = 50 mm^2^ g^−1^ min^−1^. We solve the regularised equations (2.7) on *r* ∈ [0, *R*], and *t* ∈ [0, *T*], and choose *R* = 10 and *T* = 50. Since the surface tension coefficient is difficult to determine experimentally and was assumed to be 𝒪 (1) under our scaling, we use *γ*\* = 1. All other dimensional parameters are the same as in Tam et al. [12], which provides 𝒪 (1) parameters estimated from *S. cerevisiae* mat formation experiments. The 𝒪 (1) Péclet number represents that growth-induced advection of nutrients occurs at a similar rate to diffusion. In numerical solutions, we first investigate biofilm growth assuming very strong biofilm–substratum adhesion (*λ*\* → ∞), for which the tangential slip boundary condition (2.3c) becomes the no-slip condition. Later, we will relax this assumption to investigate how *λ*\* affects solutions.

**Table 2.1:** Dimensionless parameters for the thin-film lubrication model.

| Par | Value | Par | Value | Par | Value | Par | Value |
| --- | --- | --- | --- | --- | --- | --- | --- |
| $R$ | 10 | $T$ | 50 | $h^*$ | 0.002 | $b$ | $1 \times 10^{-4}$ |
| $H_0$ | 0.1 | $\Psi_m$ | 0.111 | $\Psi_d$ | 0 | $\Upsilon$ | 3.15 |
| $D$ | 1.05 | $\text{Pe}$ | 3.94 | $Q_b$ | 8.65 | $Q_s$ | 2.09 |
| $\gamma^*$ | 1 | $\lambda^*$ | ∞ | | | | |

### 2.4 Numerical Solutions

We solve the regularised two-dimensional lubrication model (2.7) numerically using the Crank–Nicolson method. Rather than solve for *ϕ*_*n*_(*r, z, t*) directly, we instead introduce the auxiliary variable

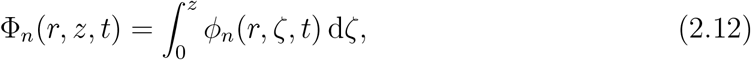

which is the cumulative cell density through a vertical slice of the biofilm, measured from the biofilm–substratum interface. We then have 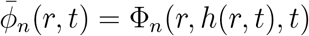, and can recover the solution for *ϕ*_*n*_ using

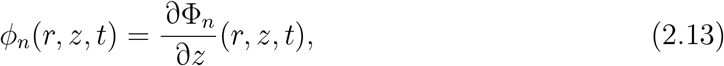

which we approximate via a centred finite-difference stencil. Integrating equation (2.7) with respect to *z* yields an equivalent equation in terms of Φ_*n*_. To simplify the spatial finite-difference stencils, we make a substitution *z* = *ξh*(*r, t*) to map Φ_*n*_(*r, ξh*(*r, t*), *t*) onto a (stationary) rectangular domain (*r, ξ*) ∈ [0, *R*] × [0, 1]. The resulting discrete system of equations is nonlinear, and we solve it using Newton’s method. We compared applying Newton’s method to the complete system of equations (2.7) simultaneously, with solving (2.7a), (2.7b), (2.7e) and (2.7f) in turn. The latter approach was more efficient, with no appreciable loss in accuracy for the range of parameters considered herein. Further details on the numerical scheme are provided in the Supplemental Material.

The numerical solution with initial conditions (2.8), boundary conditions (2.9)–(2.11), and parameters listed in Table 2.1 is shown in Figures 2.2 and 2.3. Comparing Figure 2.2 to the results of Ward and King [46] reveals how nutrient limitation and non-constant cell volume fraction affect the evolution of the biofilm. A notable change is that here we observe a non-constant expansion speed. Initially, biofilm expansion is comparatively fast because nutrients are abundant, but expansion slows as nutrients deplete. This behaviour is to be expected, because in our model the source term is proportional to *g*_*b*_, which decreases with time (see Figure 2.2d). Figure 2.2a also shows how the nutrient distribution affects biofilm shape. When nutrients are abundant, the biofilm grows both vertically and radially, and attains a thicker shape to that seen in Ward and King [46]. As nutrients deplete and cell proliferation decreases, surface tension forces facilitate slower radial spread, in conjunction with a decrease in height at *r* = 0.

**Figure 2.2:**
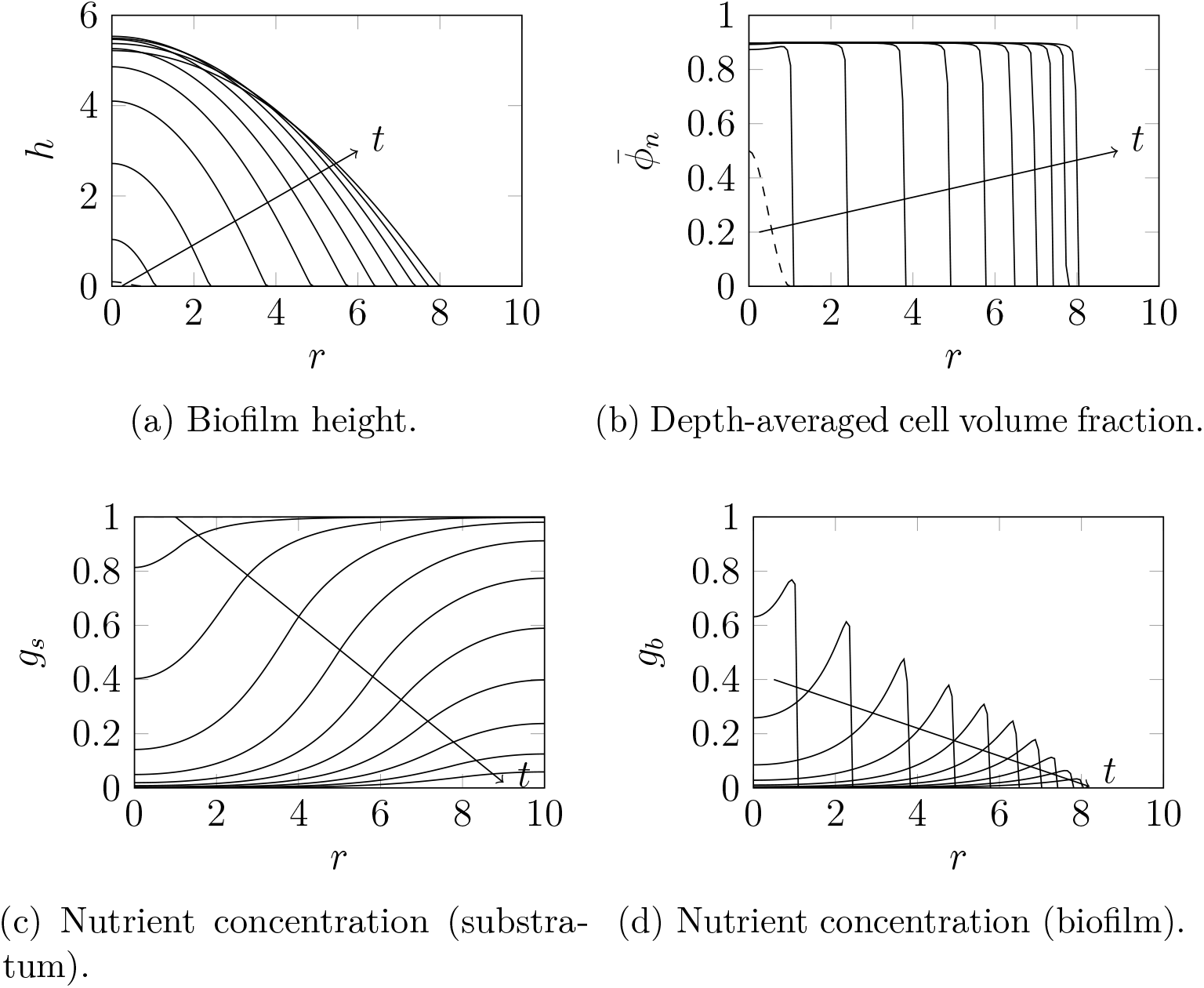
Numerical solution to (2.7), with the initial conditions (2.8) and parameters as in Table 2.1. Where visible, dashed curves represent initial conditions, and we plot solutions in increments of *t* = 5. Arrows indicate the direction of increasing time.

**Figure 2.3:**
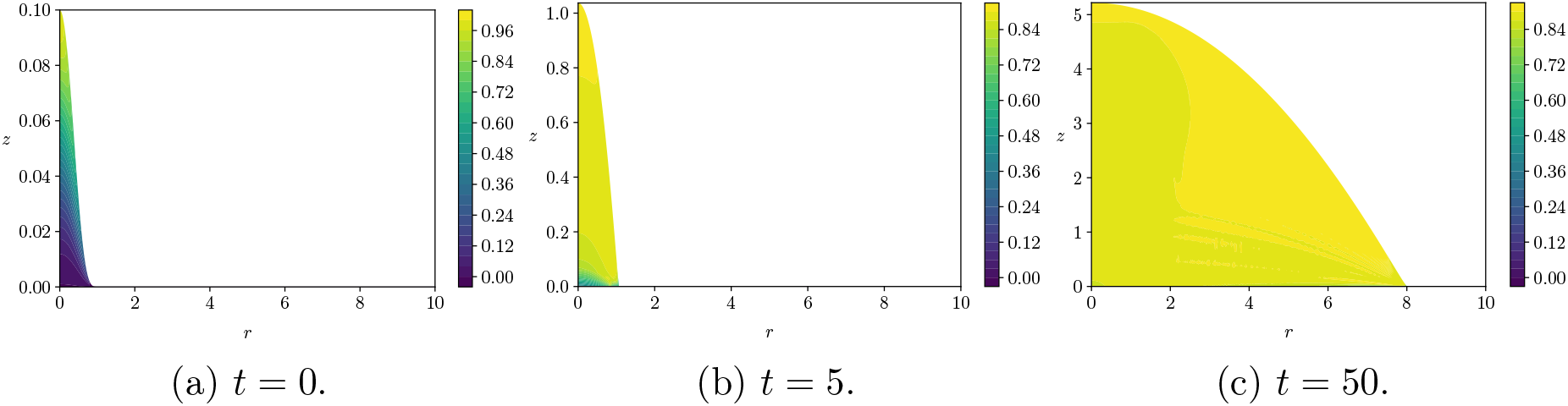
The distribution of cell volume fraction, *ϕ*_*n*_, within the biofilm, for the numerical simulations in Figure 2.2 with the initial conditions (2.8) and parameters as listed in Table 2.1.

Figure 2.3 illustrates the spatial dependence of the cell volume fraction *ϕ*_*n*_. We chose the initial condition in Figure 2.3a such that the distribution of *ϕ*_*n*_ connects continuously to the precursor film, for which *ϕ*_*n*_ = 0. During early development, we observe a region close to *z* = 0 where the cell volume fraction is low, as Figure 2.3b shows. This is a result of the initial condition (2.8b) and no-slip condition (2.3c). Over the duration of the simulation, the biofilm evolves such that the distribution of *ϕ*_*n*_ becomes close to uniform with *r* and *z*. This suggests that the initial variation and *z*-dependence in cell volume fraction has little effect on biofilm evolution. To investigate this further, we develop a simplified one-dimensional version of our model.

## 3 Simplified One-Dimensional Model

We now exploit the observation in Figure 2.3 that cell volume fraction exhibits weak dependence on *z*, and assume that *ϕ*_*n*_ = *ϕ*_*n*_(*r, t*). This enables us to reduce the two-dimensional axisymmetric model (2.7) to a one-dimensional form, and eliminate *u*_*z*_. Neglecting the two-dimensional structure has the major advantage of saving computational time. After assuming that *ϕ*_*n*_ is independent of *z*, equation (2.7c) reduces to 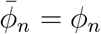, and equations (2.7a), (2.7e) and (2.7f) are unchanged. Applying the thin-film approximation yields a new equation for the cell volume fraction,

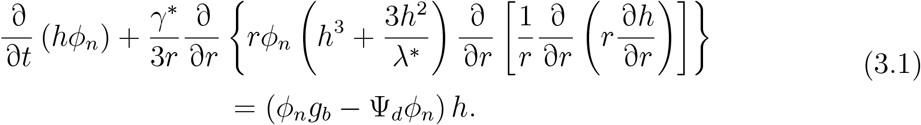

Full details on the derivation of (3.1) are provided in the Supplemental Material. Equations (3.1), (2.7a), (2.7e) and (2.7f), together with the boundary conditions (2.9)–(2.11) and initial conditions (2.8a) and (2.8d) and

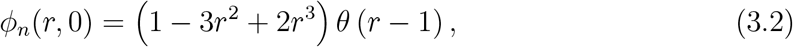

form a closed system for *h*(*r, t*), *ϕ*_*n*_(*r, t*), *g*_*s*_(*r, t*), and *g*_*b*_(*r, t*). Notably, the one-dimensional simplification eliminates the two-dimensional integro-differential equation (2.7d) for *u*_*z*_ from the thin-film model, enabling faster computation.

### 3.1 Numerical Solutions

Our numerical method for the one-dimensional model is similar to the method for the two-dimensional model described in §2.4, with the simplification that *ϕ*_*n*_(*r, t*) is now a one-dimensional field in space. As per the two-dimensional model, we introduce an auxiliary variable Φ_*n*_(*r, t*) = *h*(*r, t*)*ϕ*_*n*_(*r, t*), and obtain the governing equation for Φ_*n*_(*r, t*) from (3.1). We then apply the Crank–Nicolson method directly to the system of equations (3.1), (2.7a), (2.7e) and (2.7f), noting that 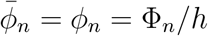. The resulting discrete system of equations is nonlinear, and we solve it using Newton’s method. Again, applying Newton’s method to each equation in turn was more efficient than applying it to all equations simultaneously, and yielded solutions with no appreciable loss of accuracy. More information on the numerical scheme is available in the Supplemental Material.

The one-dimensional numerical solution using the parameters listed in Table 2.1 is shown in Figure 3.1. The biofilm develops similarly to the two-dimensional solution in Figure 2.2. Since the mass balance equations for *h, g*_*s*_, and *g*_*b*_ are the same in both the one-dimensional and two-dimensional models, we expect these variables to develop similarly. Although the two-dimensional thin-film reduction does not allow us to assume initially that *ϕ*_*n*_ is independent of *z*, making this assumption does not significantly affect the results. Furthermore, the results in Figure 3.1b suggest that the cell volume fraction quickly becomes uniform throughout the biofilm, as we observed in Figure 2.3 for the two-dimensional model. This is consistent with the assumptions that cell volume fraction is independent of *z*, or can be assumed constant from the outset.

**Figure 3.1:**
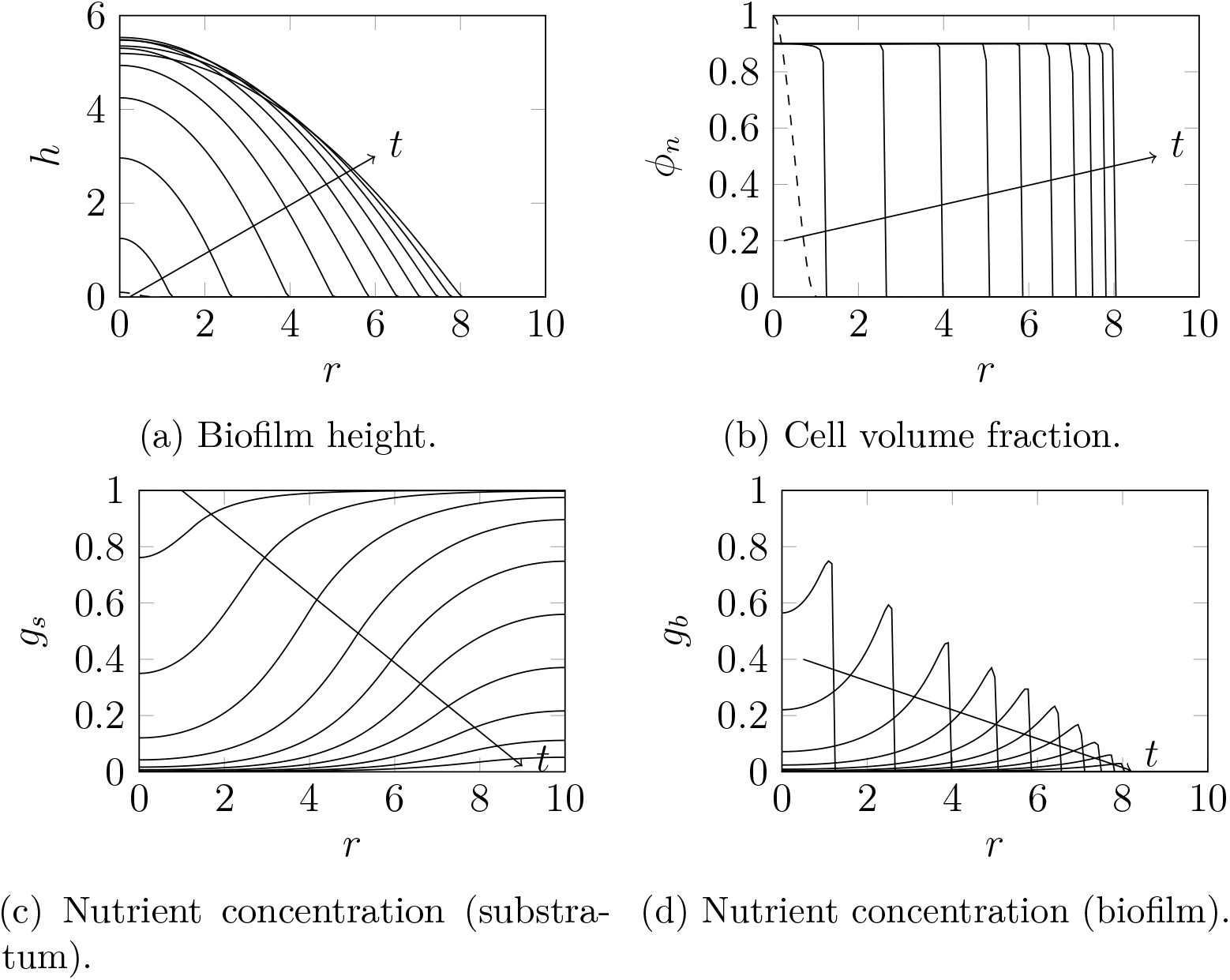
Numerical solution to the simplified one-dimensional model, with the initial conditions (3.2), (2.8a), (2.8c) and (2.8d) and parameters as in Table 2.1. Where visible, dashed curves represent initial conditions, and we plot solutions in increments of *t* = 5. Arrows indicate the direction of increasing time.

## 4 Sensitivity Analysis

To investigate the effect of parameters on biofilm expansion speed and shape, we perform a local sensitivity analysis. In each solution set, we vary one parameter from those in Table 2.1, compute a numerical solution to *t* = 25, and calculate the biofilm radius and thickness. To investigate the effect of cell proliferation rate, we vary the dimensional parameter *ψ*_*n*_, which is otherwise scaled out of the model. We then update *D*, Pe, Ψ_*m*_, and *γ*\* in each simulation based on the value of *ψ*_*n*_, and compute solutions until *T* = *ψ*_*n*_*/*2. Results for *ψ*_*n*_, Ψ_*m*_, and Ψ_*d*_ then enable us to directly compare the effects on growth of cell production rate, ECM production rate, and cell death rate respectively. Numerical results in §3.1 suggested that the simplified one-dimensional model produces qualitatively similar results to the full two-dimensional model. Performing sensitivity analyses on both models enables us to investigate this further.

Sensitivity analysis results also enable comparison between this model for strongly-adhesive biofilms and our extensional flow model [12, 55]. The extensional flow model incorporates a perfect-slip condition on the biofilm–substratum interface. This perfect-slip condition is designed to model expansion by sliding motility. Conversely, in this work the default numerical solutions for the lubrication model are for *λ*\* → ∞, representing no-slip. In the extensional flow model, we found that faster biomass production rates and increased access to nutrients at the leading edge increased biofilm expansion speed [12]. Expansion speed also depended on the initial condition, such that decreasing the initial biofilm height, *H*_0_, increased expansion speed [55]. In contrast, the surface tension coefficient, *γ*\*, has minimal effect on expansion speed in the extensional flow regime. Instead, changing *γ*\* affects biofilm shape. With low surface tension (*γ*\* < 2), the biofilm can form ridges, whereby height close to the leading edge exceeds height in the biofilm centre [12]. Here, we conduct a similar investigation for the lubrication model, to identify similarities and differences between the two regimes.

### 4.1 Biofilm Radius

One objective of the sensitivity analysis is to investigate the effect of parameters on biofilm expansion speed. The biofilm radius at *t* = 25 provides a measure of the average expansion speed in early biofilm growth. We measure the radius by computing the contact line position,

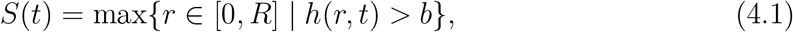

in numerical solutions. When performing the sensitivity analysis, we ensure that parameter values remain within an order of magnitude of unity, such that the lubrication scaling regime remains valid. The sensitivity analysis results, presented in Figures 4.1 and 4.2, then reveal the extent to which each parameter affects expansion speed. These results provide insight that is difficult to obtain in experiments, where physical parameters are difficult to estimate and control.

**Figure 4.1:**
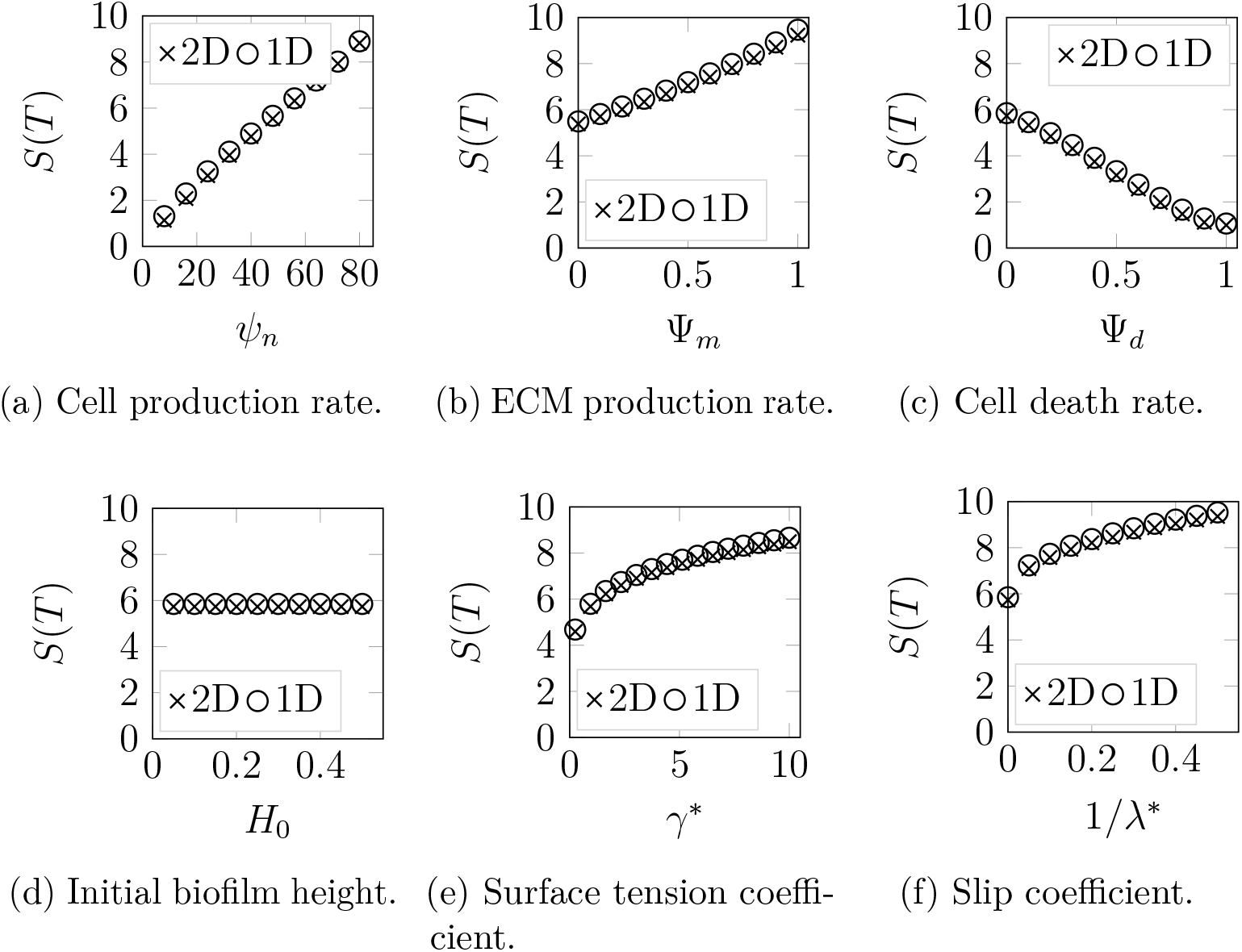
Local sensitivity analyses for biofilm radius at *t* = 25, with respect to net biomass production rates, initial biofilm height, surface tension coefficient, and slip parameter. We used the initial conditions (2.8) for all two-dimensional solutions, and replaced (2.8b) with (3.2) for one-dimensional solutions. Unless specified, all parameters used were those given in Table 2.1. Crosses represent results for the full two-dimensional model, and circles represent results for the simplified one-dimensional model.

**Figure 4.2:**
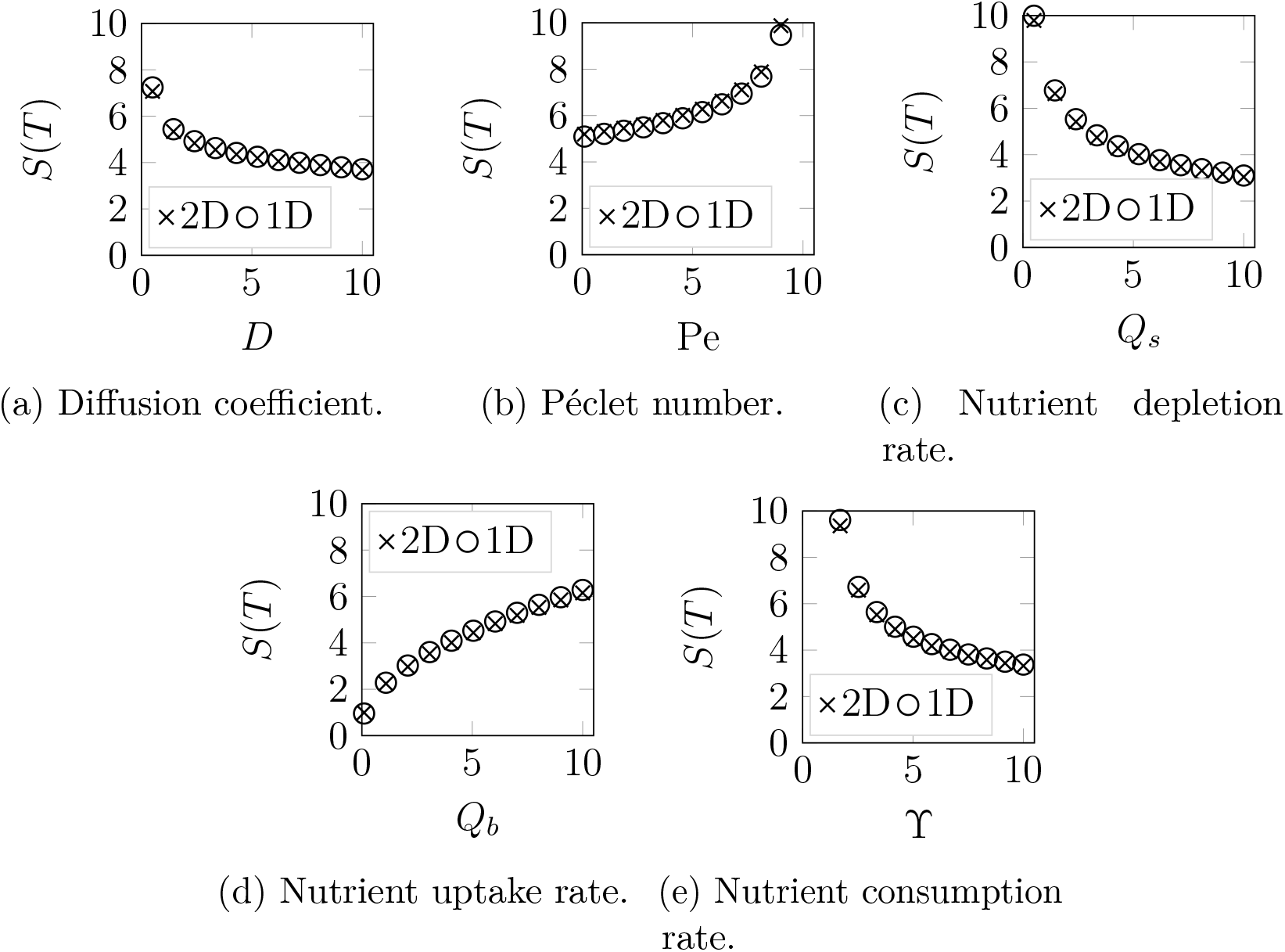
Local sensitivity analyses for biofilm radius at *t* = 25, with respect to parameters that govern the movement, consumption, and uptake of nutrients. We used the initial conditions (2.8) for all two-dimensional solutions, and replaced (2.8b) with (3.2) for one-dimensional solutions. Unless specified, all parameters used were those given in Table 2.1. Crosses represent results for the full two-dimensional model, and circles represent results for the simplified one-dimensional model.

Figure 4.1a shows that the relationship between biomass production rate and expansion speed is approximately linear, confirming that cell proliferation drives biofilm growth. As expected, increased production of extracellular fluid increases biofilm volume and subsequently expansion speed, but increased cell death rate decreases the number of cells available to proliferate, and therefore slows growth (see Figures 4.1b and 4.1c). Whilst the trends for *ψ*_*n*_, Ψ_*m*_, and Ψ_*d*_ are similar to the extensional flow results of Tam et al. [12], the effect of *γ*\* differs in the two regimes. In the lubrication model presented here, there is competition between strong biofilm–substratum adhesion and surface tension effects. When cells proliferate, strong adhesion opposes radial expansion, which restricts biofilm growth to the vertical direction. Conversely, surface tension forces oppose curvature on the free surface, and this curvature increases when the biofilm grows vertically. Surface tension forces subsequently flatten the biofilm profile and transport mass radially, facilitating biofilm expansion. We therefore observe faster radial expansion when the surface tension coefficient, *γ*\*, is increased.

The difference in expansion mechanisms between the lubrication and extensional flow regimes also explains the observation in Figure 4.1d that initial biofilm height *H*_0_ has negligible effect on biofilm radius. In the extensional flow regime, thinner biofilms expand quickly, because a relatively smaller quantity of new cells at the leading edge is required for the biofilm to spread [55]. However, due to the strong biofilm–substratum adhesion, this mechanism is not available in the lubrication regime, where instead surface tension forces determine biofilm thickness. When the other parameters are held constant, the biofilm will tend to adopt the same final shape regardless of initial height, and expand at the same speed. This also explains why the initial condition does not affect biofilm thickness in our lubrication model, as Figure 4.3d will show. Ward and King [46] observe the same behaviour in their model, whereby expansion depends on the initial conditions in the extensional flow regime, but does not in the lubrication regime. All of these conclusions apply to the case for *λ*\* → ∞, for which there is no slip at the biofilm–substratum boundary. As the resistance to slip *λ*\* decreases (*i*.*e*. 1*/λ*\* increases), we observe faster expansion (see Figure 4.1f). This is because relaxing the *λ*\* → ∞ assumption introduces slip, enabling the biofilm to spread as cells proliferate at the leading edge. This spread occurs without the need for surface tension forces to redistribute mass radially, giving rise to non-zero radial velocity at the contact line. It also enables the biofilm to invade potentially more nutrient-rich regions of the Petri dish closer to its edge.

**Figure 4.3:**
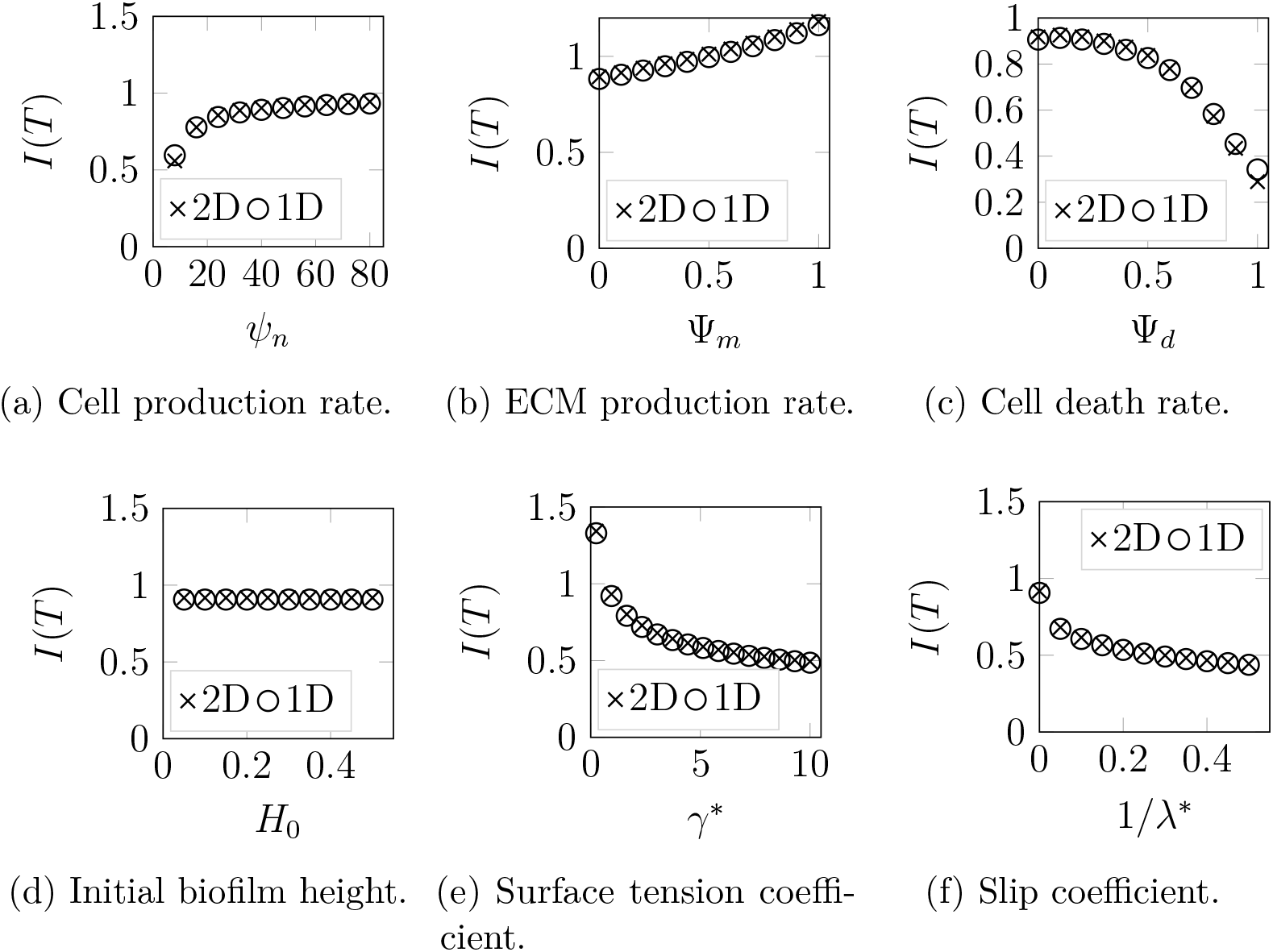
Local sensitivity analyses for biofilm aspect ratio at *t* = 25, with respect to net biomass production rates, initial biofilm height, surface tension coefficient, and slip parameter. We used the initial conditions (2.8) for all two-dimensional solutions, and replaced (2.8b) with (3.2) for one-dimensional solutions. Unless specified, all parameters used were those given in Table 2.1. Crosses represent results for the full two-dimensional model, and circles represent results for the simplified one-dimensional model.

The effects of parameters associated with nutrients on expansion speed are shown in Figure 4.2. Increasing the nutrient diffusion coefficient, *D*, will enable faster movement of nutrients towards the centre of the biofilm, replenishing consumed nutrients and resulting in more uniform nutrient concentrations across the Petri dish. This promotes thickening of the biofilm as opposed to radial expansion. Increasing the nutrient consumption rate,ϒ, has the opposite effect of slowing expansion, because it results in larger quantities of nutrient being required to produce a new cell. The Péclet number indicates how readily nutrients advect radially with the extracellular fluid. Larger values of Pe increase nutrient supply to the proliferating rim, enabling faster expansion. Since biofilms tend to be thicker in the lubrication regime than the extensional flow regime, advection within the biofilm has a stronger effect on nutrient availability in the lubrication regime than in the extensional flow regime. This explains the stronger effect of the Péclet number when compared to Tam et al. [12]. Larger values of nutrient depletion rate, *Q*_*s*_, decrease nutrient availability to the cells, which slows expansion. Conversely, increasing nutrient uptake rate, *Q*_*b*_, aids cell production, because more nutrients become available for consumption.

In summary, expansion speed in strongly-adhesive biofilms depends on a combination of cell proliferation (mediated by nutrient availability), surface tension forces, and slip. Parameter changes that increase cell proliferation cause faster expansion, because they increase the quantity of biomass. In contrast, increased surface tension forces, which represents increased cell–cell adhesion on the upper surface of the biofilm, redistribute biomass radially. This generates faster expansion, but also leads to thinner biofilms. Similarly, the addition of slip on the biofilm–substratum interface promotes radial expansion at the expense of vertical growth, giving rise to faster expansion. We reinforce these conclusions in §4.2, where we investigate the effect of parameters on biofilm thickness.

### 4.2 Biofilm Thickness

Computing the biofilm thickness in the sensitivity solutions provides a way of quantifying the effect of parameters on biofilm shape. For each solution in the sensitivity analysis, we compute the dimensionless aspect ratio,

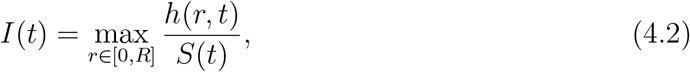

to measure the relative biofilm thickness. Sensitivity analysis results for this aspect ratio are presented in Figures 4.3 and 4.4.

**Figure 4.4:**
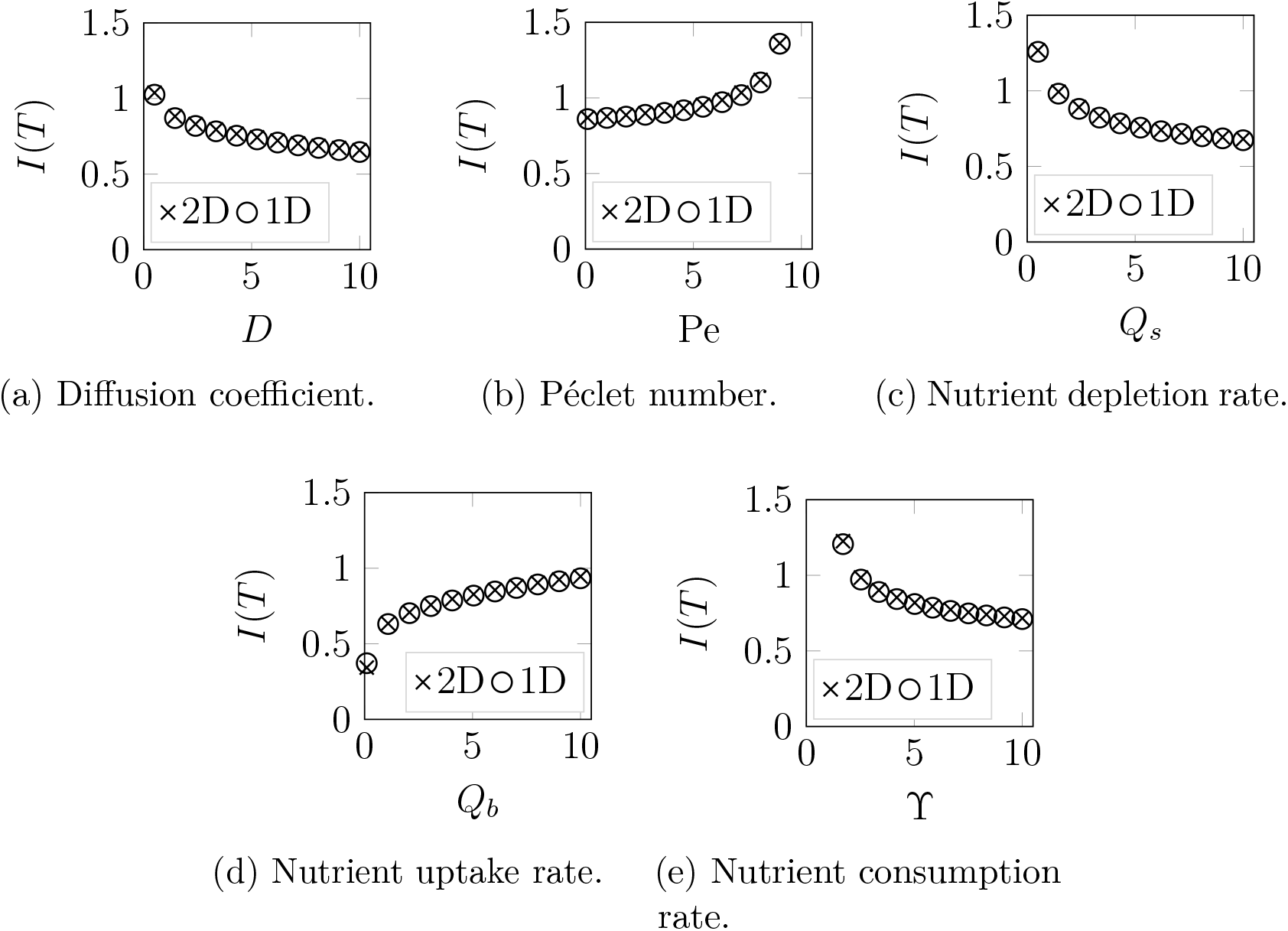
Local sensitivity analyses for biofilm aspect ratio at *t* = 25, with respect to parameters that govern the movement, consumption, and uptake of nutrients. We used the initial conditions (2.8) for all two-dimensional solutions, and replaced (2.8b) with (3.2) for one-dimensional solutions. Unless specified, all parameters used were those given in Table 2.1. Crosses represent results for the full two-dimensional model, and circles represent results for the simplified one-dimensional model.

As described in §4.1, Figure 4.3d shows that the initial biofilm height has negligible impact on thickness. Most of the parameters that affect the biofilm thickness, including Ψ_*m*_, Ψ_*d*_, and all parameters in Figure 4.4, do so because they affect the quantity of biomass. Increasing the quantity of biomass promotes both radial growth and thickening of the biofilm, and therefore changes that increase expansion speed also increase biofilm thickness. In contrast, the surface tension coefficient *γ*\* and the slip parameter *λ*\* affect the distribution of biomass in the biofilm, and not its quantity. Since increasing *γ*\* increased the radial size, this occurs in conjunction with a reduction in thickness, as Figure 4.3e shows. Similarly, decreasing the resistance to slip promotes radial expansion as opposed to thickening, and therefore thickness increases as 1*/λ*\* increases.

In all solutions presented in Figures 4.3 and 4.4, the biofilm aspect ratio remains close to unity. By contrast, biofilms in the extensional flow regime are thinner, with *I* < 0.05 observed in the solutions of Tam et al. [12]. This is because the extensional flow model incorporates perfect-slip on the biofilm–substratum interface, facilitating radial spread as cells proliferate. In contrast, *γ*\* mediates radial spread in the lubrication regime, and biofilms with *I* ≈ 1 emerge with *γ*\* = 1. These results suggest that biofilm aspect ratio provides a means to distinguish between the extensional flow and lubrication regimes, and subsequently the dominant expansion mechanism. For example, *S. cerevisiae* mat biofilms [10, 11] and *B. subtilis* biofilms [56] can grow with thin aspect ratios of *I* ≈ 0.02, suggesting that sliding motility governs their growth. In addition, numerical solutions can contain ridges in the extensional flow regime [12], but not in the lubrication regime. Therefore, the presence of ridged colony shape, for example in the experiments of Maršíková et al. [57], might also indicate expansion governed by sliding motility, and not strong adhesion.

Throughout Figures 4.3 and 4.4, we observe similar trends in biofilm thickness between the two-dimensional and one-dimensional models. Combining this with the results of Figures 4.1 and 4.2, we conclude that the one-dimensional model captures the key mechanisms of the lubrication regime, including the effects of cell proliferation, nutrient transport and uptake, and surface tension. Therefore, although the one-dimensional model does not capture the vertical variation in the cell volume fraction, it provides a viable alternative to the two-dimensional model for the biologically-feasible parameters considered here. The advantage of the one-dimensional model is that it is computationally less expensive, and potentially more amenable to analysis, than the two-dimensional model.

## 5 Discussion and Conclusion

We have derived a mathematical model for the growth of biofilms that adhere strongly to a surface. The model describes biofilms as two-phase fluids, consisting of living cells and extracellular fluid. Our model extends the thin-film lubrication model of Ward and King [46] beyond the assumption of early biofilm growth. To achieve this, we allow the volume fractions of living cells and extracellular material to vary throughout the biofilm, and explicitly model nutrient uptake and depletion. After deriving the model, we computed radially-symmetric numerical solutions to the full model, and a one-dimensional simplification that neglected vertical variation in volume fraction. Finally, we performed a local sensitivity analysis, by varying each parameter and investigating its effect on biofilm radius and thickness. We found that increased production of cells and extracellular material, increased nutrient availability, increased surface tension, and increased slip on the biofilm–substratum interface all facilitate faster biofilm growth.

To our knowledge, this work is the first to investigate the vertical dependence of volume fraction within a multi-phase, thin-film model for biofilm growth. Previous biofilm models (see [32, 37, 41, 45–48]) assumed that the volume fraction of active biomass is constant or independent of *z*. In our model, this assumption cannot be made *a priori*. However, we found good agreement between solutions to the full model and a simplified one-dimensional model that neglects possible *z*-dependence in volume fractions. Since our results show that models neglecting *z*-independence can capture the important mechanisms of biofilm growth, our study reinforces the conclusions of prior models.

The parameter sensitivity analysis identified qualitative differences between results in the lubrication regime considered here, and the extensional flow regime considered in Tam et al. [12]. In the lubrication regime, expansion speed depends on surface tension coefficient, *γ*\*, which has negligible effect on speed in the extensional flow regime. Surfactants have been shown to disrupt bacterial biofilm growth [58, 59]. The effect of surfactants could be used to identify the dominant mechanisms of growth in an experiment. If the surfactant inhibits biofilm growth, then our results suggest the lubrication regime applies, with biofilm growth driven by surface tension as cells proliferate. The biofilm aspect ratio provides another means of distinguishing between the two regimes. Our results suggest that sliding motility gives rise to thin biofilms, whereas biofilms driven by surface tension and strong adhesion will has aspect ratios closer to unity. This can be investigated in an experiment by observation, or by measuring the biofilm height, for example using an electrometer [60]. Furthermore, we found that ridge formation only occurs in the extensional flow regime, which would suggest expansion by sliding motility. Comparison between the sensitivity analyses of Tam et al. [12] and the present work can help experimentalists identify candidate expansion mechanisms. However, we acknowledge that these models neglect some mechanisms, for example osmotic swelling, that might also affect expansion.

If the lubrication regime is relevant, our numerical solutions and parameter sensitivity analyses provide quantitative predictions that can be tested experimentally. Nutrient diffusivity can be controlled by adjusting the weight percentage of the agar medium, according to the empirical relationship of Slade, Cremers, and Thomas [61]. The biofilm– substratum adhesion strength indicates the slip coefficient, *λ*\*, which controls whether the lubrication or extensional flow regimes is relevant. In practice, many factors affect adhesion strength [62]. For information on how to control adhesion strength in three bacterial species, we refer the interested experimentalist to the review by Jiang et al. [63]. Guidelines for measuring adhesion strength are provided in the review by Boudarel et al. [64]. This review also describes how to measure cohesion strength, which is analogous to surface tension in biofilm colonies [25]. The surface tension coefficient, *γ*\*, can be varied by application of surfactants. Knowledge of the cell proliferation and death rates is also required to compare our model with experiments. This can be attained using a red/green staining assay [65–68] to track the number of living and dead cells over time.

The research presented here contains scope for future work. Biofilms of bacteria and yeast can form a diverse range of spatiotemporal patterns, such as the floral morphology [10]. An interesting extension to this work would be to use our model to investigate these patterns by relaxing the assumption of azimuthal symmetry, and determining the stability of solutions to azimuthal perturbations. Other possible extensions include modelling a viscoelastic substratum instead of solid, or considering the general viscoelastic behaviour of the biofilm itself. One could then impose continuity of shear stress on the biofilm– substratum interface to investigate the effect of agar properties on biofilm growth. Our general model also provides a framework from which models with more detail can be developed. For example, it might be possible to observe more complicated behaviour by relaxing the assumption that the phases are mechanically equivalent, or by decomposing the biofilm into three or more phases. The model can also be extended to investigate the effects of osmotic swelling [42, 43, 47], and ECM production regulated by quorum sensing [34, 69], both of which are also hypothesised to affect biofilm growth. The addition of more complicated mechanisms might require the full two-dimensional cell volume fraction profile to accurately predict biofilm growth, unlike the solutions presented here that exhibited weak dependence on *z*.

## Acknowledgements

A. K. Y. T. acknowledges funding from the A. F. Pillow Applied Mathematics Trust, from the Australian Government under the Research Training Program, and from the Australian Research Council (ARC), under grant number DP180102956. J. E. F. G. acknowledges support from the School of Mathematical Sciences and the Faculty of Engineering, Computer and Mathematical Sciences, University of Adelaide, through the Special Studies Programme, and from the ARC (grant number DE130100031). S. B. acknowledges funding from the ARC (grant number DP200101764). B. J. B. and B. H. acknowledge funding from the ARC (grant number DP160102644).

