## Supplementary Material for "A Thin-Film Lubrication Model for Biofilm Expansion Under Strong Adhesion"

In §1 and 2, we derive the two-dimensional thin-film and simplified one-dimensional models used in the main text.

### 1 Two-Dimensional Model

To derive the two-dimensional thin-film lubrication model, we consider growth of a yeast biofilm on a solid substratum, and adopt a radially-symmetric cylindrical co-ordinate system $(r, z)$. The biofilm inhabits the region $0 < r < S(t)$, where $S(t)$ is referred to as the *contact line*. The biofilm is bounded below by a rigid substratum of thickness $H_s$, and bounded above by a free surface $z = h(r, t)$. Biofilm growth occurs with characteristic height $H_b$, and a characteristic radius $R_b$. A sketch of the problem domain, which closely resembles that of Ward and King [1], is shown in Figure 1.1.

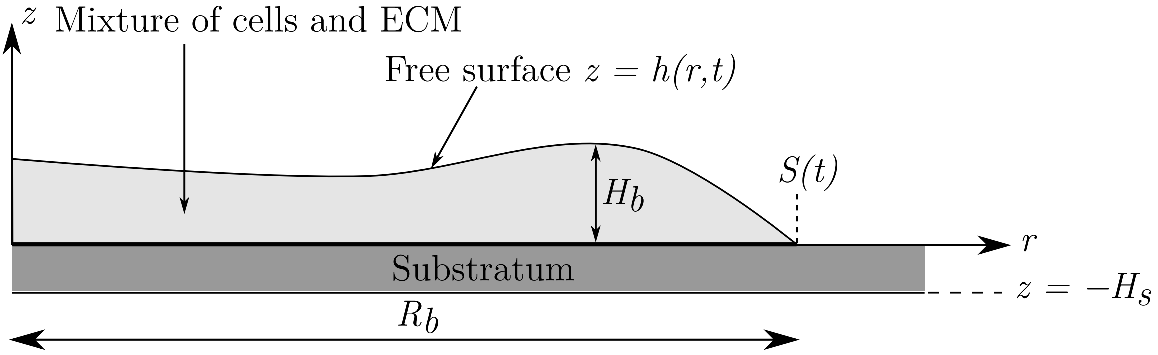


Figure 1.1: A simplified representation of a vertical slice through the centre of the biofilm and substratum. The biofilm exists in the region bounded by $0 < z < h(r, t)$, and $0 < r < S(t)$.

We adopt a macroscopic continuum modelling approach, and treat the biofilm as a mixture of two viscous fluid phases. The first phase is a living cell phase consisting of active fluid, denoted with the subscript $n$. The second phase consists of extracellular polymeric

substances and all remaining passive fluid, and we refer to this as the extracellular matrix (ECM), and denote this phase with the subscript $m$. We define the volume fractions of living cells and ECM to be $\phi_n(r, z, t)$ and $\phi_m(r, z, t)$ respectively, and assume that the fluid mixture contains no voids, that is

$$\phi_n + \phi_m = 1. \tag{1.1}$$

When defining these volume fractions we implicitly assume that an appropriate averaging process has taken place, and do not discuss the details here.

Along with biofilm mechanics, we incorporate the uptake of nutrients from the substratum. To enable this, we introduce $g_s(r, z, t)$, the nutrient concentration in the substratum defined for $-H_s < z < 0$, and $g_b(r, z, t)$, the nutrient concentration in the biofilm, defined for $0 < z < h(r, t)$ and $0 < r < S(t)$. We use two distinct nutrient concentrations because the nutrient concentration is initially discontinuous across the biofilm–substratum interface. As the biofilm evolves, nutrients can enter the biofilm across this interface, at which point they become available for consumption by the cells. We assume that nutrients disperse by diffusion in the substratum, and by both diffusion and advection with extracellular fluid inside the biofilm.

#### 1.1 Governing Equations and Boundary Conditions

We derive the governing equations of our general model using the conservation of mass and momentum for each species. For each fluid phase, we assume that the mass flux is entirely advective, and we write

$$\frac{\partial \phi_n}{\partial t} + \nabla \cdot (\phi_n \boldsymbol{u}_n) = \psi_n \phi_n g_b - \psi_d \phi_n, \tag{1.2a}$$

$$\frac{\partial \phi_m}{\partial t} + \nabla \cdot (\phi_m \boldsymbol{u}_m) = \psi_m \phi_n g_b + \psi_d \phi_n, \tag{1.2b}$$

where $\boldsymbol{u}_n$ and $\boldsymbol{u}_m$ are the fluid velocities for the living cells and ECM phases respectively. We assume linear source terms, which are the simplest forms that model fluid production proportional to local cell density and nutrient concentration. We define $\psi_n$ to be the rate of cell production, and $\psi_m$ to be the rate of ECM production, and assume both to be constant. Cell death occurs at the constant rate $\psi_d$, and dead cells immediately become part of the ECM phase with no change in volume. Since these source terms describe mass creation in the biofilm, the fluid velocities for each phase do not satisfy the usual incompressibility condition $\nabla \cdot \boldsymbol{u}_\alpha = 0$, where $\alpha = n, m$ denotes a fluid phase.

In the substratum, we assume that nutrients disperse by diffusion alone. Once nutrients enter the biofilm, they become available for consumption by the cells, and we assume that total flux consists of both diffusion and advection with the fluid phases. We also assume that the rate of nutrient consumption is proportional to the local cell volume fraction and nutrient concentration. The mass balance equations for the nutrients then read

$$\frac{\partial g_s}{\partial t} = D_s \nabla^2 g_s, \tag{1.3a}$$

$$\frac{\partial g_b}{\partial t} + \nabla \cdot \left[ g_b \left( \phi_n \boldsymbol{u}_n + \phi_m \boldsymbol{u}_m \right) \right] = D_b \nabla^2 g_b - \eta \phi_n g_b, \tag{1.3b}$$

where $D_s$ and $D_b$ are the nutrient diffusivities in the substratum and biofilm respectively, and $\eta$ is the maximum rate at which nutrient is consumed.

We obtain the remaining governing equations from the principle of momentum conservation. Since experiments show that inertial effects are negligible on the time and length scales of biofilm growth [2], the momentum balance equations for each phase are

$$\nabla \cdot (\phi_\alpha \boldsymbol{\sigma}_\alpha) + \boldsymbol{F}_\alpha = \mathbf{0}, \tag{1.4}$$

where $\boldsymbol{\sigma}$ is the stress tensor, and $\boldsymbol{F}$ represents net sources of momentum. Equation (1.4), together with the mass balance equations (1.2) and (1.3), provide the basis for our model.

To write our equations in terms of physical properties of the fluids, we require constitutive relations for the stress tensors $\boldsymbol{\sigma}_\alpha$ and momentum source terms $\boldsymbol{F}_\alpha$. These describe the mechanical behaviour of the cells and extracellular fluid. For the stress tensors, we assume that both phases are Newtonian viscous fluids. Owing to cell proliferation and local ECM production, the stress components include terms involving $\nabla \cdot \boldsymbol{u}_\alpha$, which would otherwise vanish due to incompressibility. In cylindrical co-ordinates, the relevant stress components are [3]

$$\begin{aligned} \sigma_{rr\alpha} &= -p_\alpha - \frac{2\mu_\alpha}{3} \nabla \cdot \boldsymbol{u}_\alpha, \\ \sigma_{rz\alpha} = \sigma_{zr\alpha} &= \mu_\alpha \left( \frac{\partial u_{r\alpha}}{\partial z} + \frac{\partial u_{z\alpha}}{\partial r} \right), \\ \sigma_{\theta\theta\alpha} &= -p_\alpha - \frac{2\mu_\alpha}{3} \nabla \cdot \boldsymbol{u}_\alpha + \frac{2\mu_\alpha}{r} u_{r\alpha}, \\ \sigma_{zz\alpha} &= -p_\alpha - \frac{2\mu_\alpha}{3} \nabla \cdot \boldsymbol{u}_\alpha + 2\mu_\alpha \frac{\partial u_{z\alpha}}{\partial z}, \end{aligned} \tag{1.5}$$

where for each phase $p_\alpha$ is the pressure, and $\mu_\alpha$ is the dynamic viscosity, and these viscosities for each phase are assumed constant. When writing the stress tensor components, we invoke Stokes' hypothesis, giving the coefficient $-2\mu_\alpha/3$ for the divergence terms [1, 4, 5] in (1.5). Here, we also neglect growth pressure due to cell–cell contact, which was previously considered in similar models [6, 7]. Instead, we assume that microbes cannot respond actively to chemical or mechanical cues from the environment. This is appropriate for yeasts which are non-motile, and also bacteria which often lose swimming motility in biofilm environments [4]. By making this assumption, we suggest that the incompressibility of the material is sufficient to drive expansion when cells proliferate.

Regarding the sources of momentum, we follow Green et al. [8] by assuming that the ECM exerts a drag force on the cells. We therefore prescribe the momentum sources as

$$\boldsymbol{F}_n = -k\left(\boldsymbol{u}_n - \boldsymbol{u}_m\right) + p_n \nabla\phi_n, \quad \boldsymbol{F}_m = -k\left(\boldsymbol{u}_m - \boldsymbol{u}_n\right) + p_m \nabla\phi_m, \tag{1.6}$$

where $k(\phi_n, \phi_m) \geq 0$ is the inter-phase viscous drag coefficient. The second term on the right-hand side of each momentum source (1.6) represents interfacial forces between cells and the ECM.

Now, if we substitute the constitutive relations for the stress tensors (1.5) and momentum source terms (1.6) into the momentum balance equations (1.7), we obtain

$$\begin{aligned} -\frac{\partial}{\partial r}\left(\phi_\alpha p_\alpha\right) - \frac{2\mu_\alpha}{3}\frac{\partial}{\partial r}\left(\phi_\alpha \nabla\cdot\boldsymbol{u}_\alpha\right) + \mu_\alpha \nabla\cdot\left(\phi_\alpha \frac{\partial \boldsymbol{u}_\alpha}{\partial r}\right) \\ +\mu_\alpha \nabla\cdot\left(\phi_\alpha \nabla u_{r\alpha}\right) - \frac{2\mu_\alpha\phi_\alpha}{r^2}u_{r\alpha} - k\left(u_{r\alpha} - u_{r\beta}\right) + p_\alpha \frac{\partial\phi_\alpha}{\partial r} = 0, \end{aligned} \tag{1.7a}$$

$$\begin{aligned} -\frac{\partial}{\partial z}\left(\phi_\alpha p_\alpha\right) - \frac{2\mu_\alpha}{3}\frac{\partial}{\partial z}\left(\phi_\alpha \nabla\cdot\boldsymbol{u}_\alpha\right) + \mu_\alpha \nabla\cdot\left(\phi_\alpha \frac{\partial \boldsymbol{u}_\alpha}{\partial z}\right) \\ +\mu_\alpha \nabla\cdot\left(\phi_\alpha \nabla u_{z\alpha}\right) - k\left(u_{z\alpha} - u_{z\beta}\right) + p_\alpha \frac{\partial\phi_\alpha}{\partial z} = 0, \end{aligned} \tag{1.7b}$$

where $\beta$ represents the opposite phase to $\alpha$. Given appropriate initial and boundary conditions, these momentum balance equations (1.7), together with the mass balance equations (1.2) and (1.3), define a system of governing equations for the fluid pressures, fluid velocities, and nutrient concentrations.

The first boundary condition represents that nutrient cannot pass through the base of the substratum, which is assumed to be rigid. Hence, the no-flux condition on the substratum base is

$$\left(-D_s \nabla g_s\right)\cdot\hat{\boldsymbol{n}} = \frac{\partial g_s}{\partial z} = 0, \quad \text{on} \quad z = -H_s, \tag{1.8}$$

where $\hat{\boldsymbol{n}}$ denotes the unit outward normal vector to the relevant surface. To enable cell proliferation and expansion, the biofilm takes up nutrients from the substratum. We assume that the flux of nutrients across the biofilm–substratum interface is proportional to the local concentration difference. Since there is no nutrient in the biofilm when the cells are plated, this difference is initially non-zero, and we expect that advection and consumption of nutrients in the biofilm will sustain the difference. Assuming fluid cannot pass through the interface, we then have

$$(-D_s \nabla g_s) \cdot \hat{\boldsymbol{n}} = D_s \frac{\partial g_s}{\partial z} = -Q\left(g_s - g_b\right) \quad \text{on} \quad z = 0, \tag{1.9a}$$

$$(g_b \phi_m \boldsymbol{u}_m - D_b \nabla g_b) \cdot \hat{\boldsymbol{n}} = D_b \frac{\partial g_b}{\partial z} = -Q\left(g_s - g_b\right) \quad \text{on} \quad z = 0, \tag{1.9b}$$

$$\boldsymbol{u}_\alpha \cdot \hat{\boldsymbol{n}} = u_{z\alpha} = 0 \quad \text{on} \quad z = 0, \tag{1.9c}$$

for $0 < r < S(t)$. In equations (1.9a) and (1.9b), the constant $Q$ is the nutrient mass transfer coefficient, which indicates the permeability of the biofilm. We also impose a general tangential stress condition on the substratum–biofilm interface, similar to that of Green et al. [8]. This condition reads

$$\hat{\boldsymbol{t}} \cdot (\phi_\alpha \boldsymbol{\sigma}_\alpha \cdot \hat{\boldsymbol{n}}) = -\lambda_\alpha \left(\phi_\alpha \boldsymbol{u}_\alpha \cdot \hat{\boldsymbol{t}}\right) \quad \text{on} \quad z = 0, \tag{1.10}$$

where $\hat{\boldsymbol{t}}$ is any unit tangent vector, and $\lambda_\alpha$ are coefficients representing the strength of adhesion between the fluid and substratum for each phase. This gives the general slip condition

$$\mu_\alpha \left(\frac{\partial u_{r\alpha}}{\partial z} + \frac{\partial u_{z\alpha}}{\partial r}\right) = -\lambda_\alpha u_{r\alpha}, \quad \text{on} \quad z = 0, \tag{1.11}$$

for $0 < r < S(t)$.

On the free surface, we assume that nutrient cannot pass through the biofilm–air interface. This no-flux condition is

$$(g_b \phi_m \boldsymbol{u}_m - D_b \nabla g_b) \cdot \hat{\boldsymbol{n}} = 0 \quad \text{on} \quad z = h. \tag{1.12}$$

Given that the unit outward normal to the free surface is (where the subscript $r$ denotes partial differentiation)

$$\hat{\boldsymbol{n}} = \frac{\nabla\left(z - h\right)}{\left|\nabla\left(z - h\right)\right|} = \frac{\left(-h_r, 1\right)}{\sqrt{1 + h_r^2}}, \tag{1.13}$$

this condition reads

$$g_b \phi_m \left(u_{rm} \frac{\partial h}{\partial r} - u_{zm}\right) = D_b \left(\frac{\partial g_b}{\partial r} \frac{\partial h}{\partial r} - \frac{\partial g_b}{\partial z}\right) \quad \text{on} \quad z = h. \tag{1.14}$$

We also impose the kinematic condition

$$\frac{\mathrm{D}}{\mathrm{D}t}\left(z - h\right) = \left(\frac{\partial}{\partial t} + \boldsymbol{u}_\alpha \cdot \nabla\right)\left(z - h\right) = 0, \tag{1.15}$$

on each phase, which states that fluid particles on the free surface must remain there. By expanding the material derivative and gradient operators, we can re-write this as

$$\frac{\partial h}{\partial t} + u_{r\alpha}\frac{\partial h}{\partial r} = u_{z\alpha} \quad \text{on} \quad z = h. \tag{1.16}$$

We obtain stress boundary conditions by noting that a free surface is subject to zero tangential stress, and normal stress that is proportional to its local curvature. In general, these conditions read

$$\hat{\boldsymbol{t}} \cdot (\phi_\alpha \boldsymbol{\sigma}_\alpha \cdot \hat{\boldsymbol{n}}) = 0, \quad \hat{\boldsymbol{n}} \cdot (\phi_\alpha \boldsymbol{\sigma}_\alpha \cdot \hat{\boldsymbol{n}}) = -\gamma_\alpha \kappa \quad \text{on} \quad z = h, \tag{1.17}$$

where $\gamma_\alpha$ is the surface tension coefficient of phase $\alpha$, and $\kappa = \nabla \cdot \hat{\boldsymbol{n}}$ is the mean free surface curvature. Similar to other models in biology [9], this surface tension represents the strength of cell–cell adhesion at the biofilm–air interface.

The stress tensors (1.5), and tangential and normal vectors enable us to expand the general free surface stress conditions (1.17), to obtain

$$\begin{aligned} -2\frac{\partial h}{\partial r}\left(\frac{\partial u_{r\alpha}}{\partial r} - \frac{\partial u_{z\alpha}}{\partial z}\right) + \frac{\partial u_{r\alpha}}{\partial z} + \frac{\partial u_{z\alpha}}{\partial r} \\ - \left(\frac{\partial h}{\partial r}\right)^2 \left(\frac{\partial u_{z\alpha}}{\partial r} + \frac{\partial u_{r\alpha}}{\partial z}\right) = 0 \quad \text{on} \quad z = h, \end{aligned} \tag{1.18a}$$

$$\begin{aligned} -\frac{p_\alpha}{\mu_\alpha} - \frac{2}{3}\nabla \cdot \boldsymbol{u}_\alpha + 2\left[\left(\frac{\partial h}{\partial r}\right)^2 + 1\right]^{-1} \left[\left(\frac{\partial h}{\partial r}\right)^2 \frac{\partial u_{r\alpha}}{\partial r} \right. \\ \left. - \frac{\partial h}{\partial r}\left(\frac{\partial u_{r\alpha}}{\partial z} + \frac{\partial u_{z\alpha}}{\partial r}\right)\right] = -\frac{\gamma_\alpha \kappa}{\mu_\alpha} \quad \text{on} \quad z = h, \end{aligned} \tag{1.18b}$$

where the mean curvature of the free surface is

$$\kappa = \left[\left(\frac{\partial h}{\partial r}\right)^2 + 1\right]^{-3/2} \left[-\frac{1}{r}\frac{\partial}{\partial r}\left(r\frac{\partial h}{\partial r}\right) - \frac{1}{r}\left(\frac{\partial h}{\partial r}\right)^3\right]. \tag{1.19}$$

This completes the boundary conditions associated with the model.

#### 1.2 Model Reduction

Before applying the thin-film approximation, we make further assumptions to simplify the general model. First, following O'Dea, Waters, and Byrne [10], we assume that the inter-phase drag is large, and set $k \to \infty$. Under this assumption, we need to impose that both fluid phases move with a common velocity for the momentum source terms (1.6) to remain bounded. We define this velocity to be $\boldsymbol{u} = \boldsymbol{u}_n = \boldsymbol{u}_m$. Furthermore, since both the cells and ECM are primarily composed of water, it is reasonable to expect the physical properties of each phase to be similar. We then define

$$\rho = \rho_n = \rho_m, \quad \mu = \mu_n = \mu_m, \quad \text{and} \quad \gamma = \gamma_n = \gamma_m, \tag{1.20}$$

all of which we assume constant, as well as $p = p_n = p_m$. These assumptions, combined with the no-voids assumption (1.1), reduce the governing equations (1.2), (1.3) and (1.7) to

$$\frac{1}{r}\frac{\partial}{\partial r}(ru_r) + \frac{\partial u_z}{\partial z} = (\psi_n + \psi_m)\,\phi_n g_b, \tag{1.21a}$$

$$\frac{\partial \phi_n}{\partial t} + \frac{1}{r}\frac{\partial}{\partial r}(ru_r\phi_n) + \frac{\partial}{\partial z}(u_z\phi_n) = \psi_n\phi_n g_b - \psi_d g_b, \tag{1.21b}$$

$$\frac{\partial g_s}{\partial t} = D_s\left[\frac{1}{r}\frac{\partial}{\partial r}\left(r\frac{\partial g_s}{\partial r}\right) + \frac{\partial^2 g_s}{\partial z^2}\right], \tag{1.21c}$$

$$\frac{\partial g_b}{\partial t} + \frac{1}{r}\frac{\partial}{\partial r}(ru_r g_b) + \frac{\partial}{\partial z}(u_z g_b) = D_b\left[\frac{1}{r}\frac{\partial}{\partial r}\left(r\frac{\partial g_b}{\partial r}\right) + \frac{\partial^2 g_b}{\partial z^2}\right] - \eta\phi_n g_b, \tag{1.21d}$$

$$\begin{aligned} -\frac{\partial p}{\partial r} + \frac{2\mu}{r}\frac{\partial}{\partial r}\left(r\frac{\partial u_r}{\partial r}\right) - \frac{2\mu}{3}\frac{\partial}{\partial r}\left[\frac{1}{r}\frac{\partial}{\partial r}(ru_r) + \frac{\partial u_z}{\partial z}\right] \\ +\mu\frac{\partial}{\partial z}\left(\frac{\partial u_z}{\partial r} + \frac{\partial u_r}{\partial z}\right) - \frac{2\mu}{r^2}u_r = 0, \end{aligned} \tag{1.21e}$$

$$\begin{aligned} -\frac{\partial p}{\partial z} + 2\mu\frac{\partial^2 u_z}{\partial z^2} - \frac{2\mu}{3}\frac{\partial}{\partial z}\left[\frac{1}{r}\frac{\partial}{\partial r}(ru_r) + \frac{\partial u_z}{\partial z}\right] \\ +\frac{\mu}{r}\frac{\partial}{\partial r}\left[r\left(\frac{\partial u_r}{\partial z} + \frac{\partial u_z}{\partial r}\right)\right] = 0, \end{aligned} \tag{1.21f}$$

where (1.21a) is obtained by summing the mass balance equations (1.2) for $\phi_n$ and $\phi_m$. Under these assumptions, we also have simplified forms of the boundary conditions,

$$\frac{\partial g_s}{\partial z} = 0 \quad \text{on} \quad z = -H_s, \tag{1.22a}$$

$$D_s \frac{\partial g_s}{\partial z} = -Q\left(g_s - g_b\right), \quad D_b \frac{\partial g_b}{\partial z} = -Q\left(g_s - g_b\right) \quad \text{on} \quad z = 0, \tag{1.22b}$$

$$\frac{\partial u_r}{\partial z} + \frac{\partial u_z}{\partial r} = \frac{\lambda u_r}{\mu}, \quad u_z = 0 \quad \text{on} \quad z = 0, \tag{1.22c}$$

$$g_b\left(u_r \frac{\partial h}{\partial r} - u_z\right) = D_b\left(\frac{\partial g_b}{\partial r}\frac{\partial h}{\partial r} - \frac{\partial g_b}{\partial z}\right) \quad \text{on} \quad z = h, \tag{1.22d}$$

$$\frac{\partial h}{\partial t} + u_r \frac{\partial h}{\partial r} = u_z \quad \text{on} \quad z = h, \tag{1.22e}$$

$$\begin{aligned} -2\frac{\partial h}{\partial r}\left(\frac{\partial u_r}{\partial r} - \frac{\partial u_z}{\partial z}\right) + \frac{\partial u_r}{\partial z} + \frac{\partial u_z}{\partial r} \\ -\left(\frac{\partial h}{\partial r}\right)^2\left(\frac{\partial u_z}{\partial r} + \frac{\partial u_r}{\partial z}\right) = 0 \quad \text{on} \quad z = h, \end{aligned} \tag{1.22f}$$

$$\begin{aligned} -\frac{p}{\mu} + 2\left[\left(\frac{\partial h}{\partial r}\right)^2 + 1\right]^{-1}\left[\left(\frac{\partial h}{\partial r}\right)^2 \frac{\partial u_r}{\partial r} - \frac{\partial h}{\partial r}\left(\frac{\partial u_r}{\partial z} + \frac{\partial u_z}{\partial r}\right) + \frac{\partial u_z}{\partial z}\right] \\ -\frac{2}{3}\left[\frac{1}{r}\frac{\partial}{\partial r}\left(r u_r\right) + \frac{\partial u_z}{\partial z}\right] = -\frac{\gamma\kappa}{\mu} \quad \text{on} \quad z = h. \end{aligned} \tag{1.22g}$$

We simplify the model (1.21) and (1.22) by applying the thin-film approximation. The first step in this process is to nondimensionalise the model.

#### 1.3 Scaling and Nondimensionalisation

To nondimensionalise, we first observe that the radius of a biofilm significantly exceeds both its height and the depth of the substratum. Accordingly, we assume that the aspect ratio

$$\varepsilon = \frac{H_s}{R_b} \tag{1.23}$$

is a small parameter such that $0 < \varepsilon \ll 1$, and that we also have $H_b/R_b = \mathcal{O}(\varepsilon)$ as $\varepsilon \to 0$. We then introduce the dimensionless variables denoted by hats,

$$\begin{aligned} (r, z) = (R_b\hat{r}, \varepsilon R_b\hat{z}), \quad (u_r, u_z) = (\psi_n G R_b \hat{u}_r, \varepsilon\psi_n G R_b \hat{u}_z), \\ t = \frac{\hat{t}}{\psi_n G}, \quad g_s = G\hat{g}_s, \quad g_b = G\hat{g}_b, \quad p = \frac{\psi_n G \mu}{\varepsilon^2}\hat{p}, \end{aligned} \tag{1.24}$$

where $G$ is the initial nutrient concentration. In terms of the dimensionless variables, the governing equations (1.21) become (dropping hats)

$$\frac{1}{r}\frac{\partial}{\partial r}(ru_r) + \frac{\partial u_z}{\partial z} = (1+\Psi_m)\,\phi_n g_b, \tag{1.25a}$$

$$\frac{\partial \phi_n}{\partial t} + \frac{1}{r}\frac{\partial}{\partial r}(ru_r\phi_n) + \frac{\partial}{\partial z}(u_z\phi_n) = \phi_n g_b - \Psi_d g_b, \tag{1.25b}$$

$$\frac{\partial g_s}{\partial t} = D\left[\frac{1}{r}\frac{\partial}{\partial r}\left(r\frac{\partial g_s}{\partial r}\right) + \frac{1}{\varepsilon^2}\frac{\partial^2 g_s}{\partial z^2}\right], \tag{1.25c}$$

$$\begin{aligned}\mathrm{Pe}\left[\frac{\partial g_b}{\partial t} + \frac{1}{r}\frac{\partial}{\partial r}(ru_r g_b) + \frac{\partial}{\partial z}(u_z g_b)\right] &= \frac{1}{r}\frac{\partial}{\partial r}\left(r\frac{\partial g_b}{\partial r}\right)\\ &+\frac{1}{\varepsilon^2}\frac{\partial^2 g_b}{\partial z^2} - \Upsilon\phi_n g_b,\end{aligned} \tag{1.25d}$$

$$\begin{aligned}-\frac{1}{\varepsilon^2}\frac{\partial p}{\partial r} + \frac{2}{r}\frac{\partial}{\partial r}\left(r\frac{\partial u_r}{\partial r}\right) - \frac{2}{3}\frac{\partial}{\partial r}\left[\frac{1}{r}\frac{\partial}{\partial r}(ru_r) + \frac{\partial u_z}{\partial z}\right]&\\ +\frac{\partial}{\partial z}\left(\frac{\partial u_z}{\partial r} + \frac{1}{\varepsilon^2}\frac{\partial u_r}{\partial z}\right) - \frac{2u_r}{r^2} &= 0,\end{aligned} \tag{1.25e}$$

$$\begin{aligned}-\frac{1}{\varepsilon^2}\frac{\partial p}{\partial z} + 2\frac{\partial^2 u_z}{\partial z^2} - \frac{2}{3}\frac{\partial}{\partial z}\left[\frac{1}{r}\frac{\partial}{\partial r}(ru_r) + \frac{\partial u_z}{\partial z}\right]&\\ +\frac{1}{r}\frac{\partial}{\partial r}\left[r\left(\frac{\partial u_r}{\partial z} + \varepsilon^2\frac{\partial u_z}{\partial r}\right)\right] &= 0,\end{aligned} \tag{1.25f}$$

where we have introduced the dimensionless constants

$$\begin{gathered}\Psi_m = \frac{\psi_m}{\psi_n}, \quad \Psi_d = \frac{\psi_d G}{\psi_n}, \quad D = \frac{D_s}{\psi_n G R_b^2},\\ \mathrm{Pe} = \frac{\psi_n G R_b^2}{D_b}, \quad \text{and} \quad \Upsilon = \frac{\eta R_b^2}{D_b},\end{gathered} \tag{1.26}$$

all of which we assume to be $\mathcal{O}(1)$ as $\varepsilon \to 0$. In (1.26), $\Psi_m$ and $\Psi_d$ are the dimensionless ECM production and cell death rates respectively, scaled by the cell production rate and initial nutrient concentration. The parameter $D$ is the coefficient of diffusion for nutrients in the substratum, scaled by the cell production rate and biofilm radius. The Péclet number, Pe, is the ratio of the rates of advective transport to diffusive transport within the biofilm. The parameter $\Upsilon$ is the dimensionless nutrient consumption rate. We also note that we scale $\Upsilon$ differently to the corresponding term in Ward and King [1]. In their model, the biofilm was immersed in a nutrient-rich liquid culture medium, and hence they balanced nutrient consumption with diffusion in the $z$-direction. In contrast, our biofilms grow on a nutrient-limited thin substratum, making it appropriate to balance nutrient consumption with the temporal derivative and in-plane advection and diffusion.

Applying the scaling (1.24) to the dimensionless boundary conditions, we obtain

$$\frac{\partial g_s}{\partial z} = 0 \quad \text{on} \quad z = -1, \tag{1.27a}$$

$$\frac{\partial g_s}{\partial z} = -\varepsilon^2 Q_s \left(g_s - g_b\right), \quad \frac{\partial g_b}{\partial z} = -\varepsilon^2 Q_b \left(g_s - g_b\right) \quad \text{on} \quad z = 0, \tag{1.27b}$$

$$\frac{\partial u_r}{\partial z} + \varepsilon^2 \frac{\partial u_z}{\partial r} = \lambda^* u_r, \quad u_z = 0 \quad \text{on} \quad z = 0, \tag{1.27c}$$

$$\mathrm{Pe} g_b \left(u_r \frac{\partial h}{\partial r} - u_z\right) = \frac{\partial g_b}{\partial r}\frac{\partial h}{\partial r} - \frac{1}{\varepsilon^2}\frac{\partial g_b}{\partial z} \quad \text{on} \quad z = h, \tag{1.27d}$$

$$\frac{\partial h}{\partial t} + u_r \frac{\partial h}{\partial r} = u_z \quad \text{on} \quad z = h, \tag{1.27e}$$

$$\begin{aligned} &-2\frac{\partial h}{\partial r}\left(\frac{\partial u_r}{\partial r} - \frac{\partial u_z}{\partial z}\right) + \frac{1}{\varepsilon^2}\frac{\partial u_r}{\partial z} + \frac{\partial u_z}{\partial r} \\ &-\left(\frac{\partial h}{\partial r}\right)^2 \left(\varepsilon^2 \frac{\partial u_z}{\partial r} + \frac{\partial u_r}{\partial z}\right) = 0 \quad \text{on} \quad z = h, \end{aligned} \tag{1.27f}$$

$$\begin{aligned} &-\frac{p}{\varepsilon^2} + 2\left[\varepsilon^2\left(\frac{\partial h}{\partial r}\right)^2 + 1\right]^{-1}\left[\varepsilon^2\left(\frac{\partial h}{\partial r}\right)^2 \frac{\partial u_r}{\partial r}\right. \\ &\left.-\frac{\partial h}{\partial r}\left(\frac{\partial u_r}{\partial z} + \varepsilon^2 \frac{\partial u_z}{\partial r}\right) + \frac{\partial u_z}{\partial z}\right] - \frac{2}{3}\left[\frac{1}{r}\frac{\partial}{\partial r}\left(r u_r\right) + \frac{\partial u_z}{\partial z}\right] \\ &= -\frac{\gamma^* \kappa^*}{\varepsilon^2} \quad \text{on} \quad z = h, \end{aligned} \tag{1.27g}$$

where the dimensionless parameters are

$$Q_s = \frac{Q R_b}{\varepsilon D_s}, \quad Q_b = \frac{Q R_b}{\varepsilon D_b}, \quad \lambda^* = \frac{\varepsilon \lambda R_b}{\mu}, \quad \text{and} \quad \gamma^* = \frac{\varepsilon^3 \gamma}{\psi_n G R_b \mu}. \tag{1.28}$$

Like (1.26), we also assume these to be $\mathcal{O}(1)$ as $\varepsilon \to 0$. In (1.28), $Q_s$ is a coefficient that describes the rate of nutrient depletion in the substratum, and $Q_b$ describes the rate of nutrient uptake by the biofilm. The parameter $\lambda^*$ is a dimensionless slip coefficient, and $\gamma^*$ (the reciprocal of the capillary number) represents the ratio of surface tension to viscous forces. The normal stress condition (1.27g) also depends on the dimensionless free surface curvature, $\kappa^*$, which is given by

$$\kappa^* = \left[\varepsilon^2\left(\frac{\partial h}{\partial r}\right)^2 + 1\right]^{-3/2}\left[-\frac{1}{r}\frac{\partial}{\partial r}\left(r\frac{\partial h}{\partial r}\right) - \frac{\varepsilon^2}{r}\left(\frac{\partial h}{\partial r}\right)^3\right]. \tag{1.29}$$

The governing equations (1.25) and boundary conditions (1.27) then complete the dimensionless form of our extensional flow model, on which we apply the thin-film reduction.

#### 1.4 Thin-Film Equations

The thin-film assumption introduced in §1.3 enables us to systematically simplify the dimensionless model (1.25) and (1.27). We achieve this by expanding the dependent variables as asymptotic series in powers of $\varepsilon^2$,

$$h(r,t) \sim h_0(r,t) + \varepsilon^2 h_1(r,t) + \mathcal{O}\left(\varepsilon^4\right), \tag{1.30a}$$

$$\phi_n(r,z,t) \sim \phi_{n0}(r,z,t) + \varepsilon^2 \phi_{n1}(r,z,t) + \mathcal{O}\left(\varepsilon^4\right), \tag{1.30b}$$

and so on, where series for $p$, $u_r$, $u_z$, $g_s$, and $g_b$ take the same form as (1.30b). Substituting the expansions (1.30) into the governing equations (1.25) and boundary conditions (1.27) enables us to balance physical effects of similar magnitude. In practice, we simplify the model by considering the leading-order behaviour only, which represents the strongest physical features, the remainder being $\mathcal{O}(\varepsilon^2)$ as $\varepsilon \to 0$, and hence much less significant. Applying this process to our fluid model, at leading order we obtain

$$\frac{1}{r}\frac{\partial}{\partial r}\left(r u_{r0}\right) + \frac{\partial u_{z0}}{\partial z} = \left(1 + \Psi_m\right)\phi_{n0} g_{b0}, \tag{1.31a}$$

$$\frac{\partial \phi_{n0}}{\partial t} + \frac{1}{r}\frac{\partial}{\partial r}\left(r \phi_{n0} u_{r0}\right) + \frac{\partial}{\partial z}\left(\phi_{n0} u_{z0}\right) = \phi_{n0} g_{b0} - \Psi_d \phi_{n0}, \tag{1.31b}$$

$$\frac{\partial^2 g_{s0}}{\partial z^2} = \frac{\partial^2 g_{b0}}{\partial z^2} = 0, \tag{1.31c}$$

$$\frac{\partial p_0}{\partial r} = \frac{\partial^2 u_{r0}}{\partial z^2}, \tag{1.31d}$$

$$\frac{\partial p_0}{\partial z} = 0. \tag{1.31e}$$

The leading-order boundary conditions are

$$\frac{\partial g_{s0}}{\partial z} = 0 \quad \text{on} \quad z = -1, 0, \tag{1.32a}$$

$$\frac{\partial g_{b0}}{\partial z} = 0 \quad \text{on} \quad z = 0, h_0, \tag{1.32b}$$

$$\frac{\partial u_{r0}}{\partial z} = \lambda^* u_{r0}, \quad u_{z0} = 0 \quad \text{on} \quad z = 0, \tag{1.32c}$$

$$\frac{\partial u_{r0}}{\partial z} = 0 \quad \text{on} \quad z = h_0, \tag{1.32d}$$

$$\frac{\partial h_0}{\partial t} + u_{r0}\frac{\partial h_0}{\partial r} = u_{z0} \quad \text{on} \quad z = h_0, \tag{1.32e}$$

$$p_0 = -\frac{\gamma^*}{r}\frac{\partial}{\partial r}\left(r\frac{\partial h_0}{\partial r}\right) \quad \text{on} \quad z = h_0. \tag{1.32f}$$

In (1.32f), the right-hand side term involves the leading-order contribution of the dimensionless free surface curvature, $\kappa^* = \nabla^2 h_0$.

The next step is to derive a closed system of equations in terms of leading-order quantities. First, we integrate (1.31e) with respect to $z$ and apply the normal stress boundary condition (1.32f) to obtain

$$p_0 = -\frac{\gamma^*}{r}\frac{\partial}{\partial r}\left(r\frac{\partial h_0}{\partial r}\right). \tag{1.33}$$

By (1.33) and (1.32f), the pressure throughout the biofilm is equal to the pressure on the free surface. We can use this to obtain explicit formulae for the leading-order fluid velocity components. Integrating the leading-order radial momentum equation (1.31d) twice with respect to $z$, and applying the conditions (1.32c) and (1.32d), we obtain

$$u_{r0} = \left(\frac{z^2}{2} - zh_0 - \frac{h_0}{\lambda^*}\right)\frac{\partial p_0}{\partial r}. \tag{1.34}$$

Using (1.33) to eliminate the pressure then gives

$$u_{r0} = -\gamma^*\left(\frac{z^2}{2} - zh_0 - \frac{h_0}{\lambda^*}\right)\frac{\partial}{\partial r}\left[\frac{1}{r}\frac{\partial}{\partial r}\left(r\frac{\partial h_0}{\partial r}\right)\right]. \tag{1.35}$$

We now derive leading-order mass conservation equations for the fluids in terms of the biofilm height. Integrating (1.31a) with respect to $z$ across the biofilm depth yields, on application of the kinematic (1.32e) and no-penetration (1.32c) conditions,

$$\frac{\partial h_0}{\partial t} + u_{r0}|_{z=h_0}\frac{\partial h_0}{\partial r} = (1+\Psi_m)\,\bar{\phi}_{n0}g_{b0}h_0 - \frac{1}{r}\int_0^{h_0}\frac{\partial}{\partial r}(ru_{r0})\,\mathrm{d}z, \tag{1.36}$$

where

$$\bar{\phi}_{n0} = \frac{1}{h}\int_0^{h_0}\phi_{n0}\,\mathrm{d}z. \tag{1.37}$$

To evaluate the right-hand side of (1.36), we use Leibniz's integral rule to obtain

$$\begin{aligned}\frac{\partial h_0}{\partial t} + u_{r0}|_{z=h_0}\frac{\partial h_0}{\partial r} &= (1+\Psi_m)\,\bar{\phi}_{n0}g_{b0}h_0\\ -\frac{1}{r}&\left[\frac{\partial}{\partial r}\left(\int_0^{h_0} ru_{r0}\,\mathrm{d}z\right) - ru_{r0}|_{z=h_0}\frac{\partial h_0}{\partial r}\right].\end{aligned} \tag{1.38}$$

All terms evaluated at the free surface then cancel, yielding

$$\frac{\partial h_0}{\partial t} = (1+\Psi_m)\,\bar{\phi}_{n0}g_{b0}h_0 - \frac{1}{r}\frac{\partial}{\partial r}\left(\int_0^{h_0} ru_{r0}\,\mathrm{d}z\right). \tag{1.39}$$

On replacing the velocity terms in (1.39) with the explicit formula (1.35), we obtain the leading-order conservation of total fluid mass equation,

$$\frac{\partial h_0}{\partial t} + \frac{\gamma^*}{3r}\frac{\partial}{\partial r}\left\{r\left(h_0^3 + \frac{3h_0^2}{\lambda^*}\right)\frac{\partial}{\partial r}\left[\frac{1}{r}\frac{\partial}{\partial r}\left(r\frac{\partial h_0}{\partial r}\right)\right]\right\} = (1+\Psi_m)\,\bar{\phi}_{n0}g_{b0}h_0. \tag{1.40}$$

Since the leading-order cell volume fraction $\phi_{n0}$ and the fluid velocity components (1.35) both depend on $z$, a similar approach based on vertical integration is not possible for the cellular phase mass conservation equation. Instead, we retain the three-dimensional mass balance equation (1.31b). Substituting the known radial velocity (1.35) into (1.31b), we obtain

$$\frac{\partial \phi_{n0}}{\partial t} - \frac{\gamma^*}{r}\frac{\partial}{\partial r}\left\{r\phi_{n0}\left(\frac{z^2}{2} - zh_0 - \frac{h_0}{\lambda^*}\right)\frac{\partial}{\partial r}\left[\frac{1}{r}\frac{\partial}{\partial r}\left(r\frac{\partial h_0}{\partial r}\right)\right]\right\} + \frac{\partial}{\partial z}\left(u_{z0}\phi_{n0}\right) = \phi_{n0}g_{b0} - \Psi_d\phi_{n0}. \tag{1.41}$$

Since we cannot integrate out the $z$ dependence, solving the model requires keeping track of $u_{z0}$. We achieve this by integrating (1.31a) with respect to $z$, and applying the no-penetration boundary condition (1.32c), which gives

$$u_{z0} = (1+\Psi_m)\,g_{b0}\int_0^z \phi_{n0}\,\mathrm{d}\tilde{z} + \gamma^*\int_0^z \frac{1}{r}\frac{\partial}{\partial r}\left\{r\left(\frac{\tilde{z}^2}{2} - \tilde{z}h_0 - \frac{h_0}{\lambda^*}\right)\frac{\partial}{\partial r}\left[\frac{1}{r}\frac{\partial}{\partial r}\left(r\frac{\partial h_0}{\partial r}\right)\right]\right\}\mathrm{d}\tilde{z} \tag{1.42}$$

Evaluating the second integral in (1.42) analytically, we obtain

$$u_{z0} = (1+\Psi_m)\,g_{b0}\int_0^z \phi_{n0}\,\mathrm{d}\tilde{z} + \frac{\gamma^*}{r}\frac{\partial}{\partial r}\left\{rz\left(\frac{z^2}{6} - \frac{zh_0}{2} - \frac{h_0}{\lambda^*}\right)\frac{\partial}{\partial r}\left[\frac{1}{r}\frac{\partial}{\partial r}\left(r\frac{\partial h_0}{\partial r}\right)\right]\right\}, \tag{1.43}$$

which is our leading-order equation for the vertical component of fluid velocity.

To close the model, we consider higher-order correction terms to the governing equations (1.25c) and (1.25d) to derive leading-order equations for the nutrient concentrations. Upon substituting the expansions (1.30), matching $\mathcal{O}(1)$ terms gives

$$\frac{\partial^2 g_{s1}}{\partial z^2} = \frac{1}{D}\frac{\partial g_{s0}}{\partial t} - \frac{1}{r}\frac{\partial}{\partial r}\left(r\frac{\partial g_{s0}}{\partial r}\right), \tag{1.44a}$$

$$\frac{\partial^2 g_{b1}}{\partial z^2} = \mathrm{Pe}\left[\frac{\partial g_{b0}}{\partial t} + \frac{1}{r}\frac{\partial}{\partial r}\left(ru_{r0}g_{b0}\right) + \frac{\partial}{\partial z}\left(u_{z0}g_{b0}\right)\right] - \frac{1}{r}\frac{\partial}{\partial r}\left(r\frac{\partial g_{b0}}{\partial r}\right) + \Upsilon\phi_{n0}g_{b0}. \tag{1.44b}$$

Using (1.27a), (1.27b) and (1.27d), we obtain the higher-order corrections to the boundary conditions,

$$\frac{\partial g_{s1}}{\partial z} = 0 \quad \text{on} \quad z = -1, \tag{1.45a}$$

$$\frac{\partial g_{s1}}{\partial z} = -Q_s \left(g_{s0} - g_{b0}\right) \quad \text{on} \quad z = 0, \tag{1.45b}$$

$$\frac{\partial g_{b1}}{\partial z} = -Q_b \left(g_{s0} - g_{b0}\right) \quad \text{on} \quad z = 0, \tag{1.45c}$$

$$\frac{\partial g_{b1}}{\partial z} = -\mathrm{Pe} g_{b0} \left(u_{r0}\frac{\partial h_0}{\partial r} - u_{z0}\right) + \frac{\partial g_{b0}}{\partial r}\frac{\partial h_0}{\partial r} \quad \text{on} \quad z = h_0. \tag{1.45d}$$

To derive an equation for the nutrient concentration in the substratum, we integrate (1.44a) with respect to $z$ across the substratum depth, which gives

$$\left[\frac{\partial g_{s1}}{\partial z}\right]_{-1}^{0} = \frac{1}{D}\frac{\partial g_{s0}}{\partial t} - \frac{1}{r}\frac{\partial}{\partial r}\left(r\frac{\partial g_{s0}}{\partial r}\right). \tag{1.46}$$

On applying the boundary conditions (1.45a) and (1.45b), we obtain the $z$-independent leading-order mass balance equations for nutrients in the substratum,

$$\frac{\partial g_{s0}}{\partial t} = \frac{D}{r}\frac{\partial}{\partial r}\left(r\frac{\partial g_{s0}}{\partial r}\right) - DQ_s \left(g_{s0} - g_{b0}\right), \tag{1.47}$$

Similarly, to obtain an equation for $g_{b0}$, we integrate (1.44b) with respect to $z$ across the biofilm depth to obtain

$$\begin{aligned}\mathrm{Pe}\left[h_0\frac{\partial g_{b0}}{\partial t} + \frac{1}{r}\frac{\partial}{\partial r}\left(r g_{b0}\int_0^{h_0} u_{r0}\,\mathrm{d}z\right)\right] &= \frac{1}{r}\frac{\partial}{\partial r}\left(r h_0 \frac{\partial g_{b0}}{\partial r}\right) \\ &+Q_b\left(g_{s0} - g_{b0}\right) - \Upsilon\bar{\phi}_{n0} g_{b0} h_0.\end{aligned} \tag{1.48}$$

Substituting the leading-order radial velocity (1.35) into (1.48) then yields

$$\begin{aligned}&\mathrm{Pe} h_0\frac{\partial g_{b0}}{\partial t} + \frac{\mathrm{Pe}\gamma^*}{3r}\frac{\partial}{\partial r}\left\{r g_{b0}\left(h_0^3 + \frac{3h_0^2}{\lambda^*}\right)\frac{\partial}{\partial r}\left[\frac{1}{r}\frac{\partial}{\partial r}\left(r\frac{\partial h_0}{\partial r}\right)\right]\right\} \\ &= \frac{1}{r}\frac{\partial}{\partial r}\left(r h_0\frac{\partial g_{b0}}{\partial r}\right) + Q_b\left(g_{s0} - g_{b0}\right) - \Upsilon\bar{\phi}_{n0} g_{b0} h_0,\end{aligned} \tag{1.49}$$

Equations (1.37), (1.40), (1.41), (1.43), (1.47) and (1.49) now form a closed system of equations for $h_0$, $\phi_{n0}$, $\bar{\phi}_{n0}$, $u_{z0}$, $g_{s0}$ and $g_{b0}$. These equations constitute our two-dimensional thin-film model.

### 2 One-Dimensional Model

To derive the simplified one-dimensional model, we assume that the cell volume fraction $\phi_n(r,t)$ is independent of $z$. We then obtain a new equation for $\phi_n$ by integrating the leading-order mass balance equation for living cells. After applying the boundary conditions, we obtain

$$\frac{\partial}{\partial t}(h\phi_n) + \frac{\gamma^*}{3r}\frac{\partial}{\partial r}\left\{r\phi_n\left(h^3+\frac{3h^2}{\lambda^*}\right)\frac{\partial}{\partial r}\left[\frac{1}{r}\frac{\partial}{\partial r}\left(r\frac{\partial h}{\partial r}\right)\right]\right\} \\ = \left(\phi_n g_b - \Psi_d\phi_n\right)h. \tag{2.1}$$

Here, we can use the total fluid mass conservation equation (1.40) to simplify (2.1). Multiplying (1.40) by $\phi_n$, we can then subtract the result from (2.1), and divide by $h$ to obtain

$$\frac{\partial \phi_n}{\partial t} + \frac{\gamma^*}{3}\left(h^2+\frac{3h}{\lambda^*}\right)\frac{\partial}{\partial r}\left[\frac{1}{r}\frac{\partial}{\partial r}\left(r\frac{\partial h}{\partial r}\right)\right]\frac{\partial \phi_n}{\partial r} \\ = \phi_n\left[g_b - \Psi_d - \left(1+\Psi_m\right)\phi_n g_b\right]. \tag{2.2}$$

This completes the derivation of the simplified one-dimensional model used in the main text.

### 3 Numerical Methods

Here, we describe the numerical methods used to solve both the full two-dimensional and simplified one-dimensional thin-film models. We begin by considering the two-dimensional regularised model, which is

$$\frac{\partial h}{\partial t}+\frac{\gamma^*}{3r}\frac{\partial}{\partial r}\left\{r\left(h^3+\frac{3h^2}{\lambda^*}\right)\frac{\partial}{\partial r}\left[\frac{1}{r}\frac{\partial}{\partial r}\left(r\frac{\partial h}{\partial r}\right)\right]\right\} \\ =\theta\left(h-h^*\right)\left[\left(1+\Psi_m\right)\bar{\phi}_n g_b h\right], \tag{3.1a}$$

$$\frac{\partial \phi_n}{\partial t}-\frac{\gamma^*}{r}\frac{\partial}{\partial r}\left\{r\phi_n\left(\frac{z^2}{2}-zh-\frac{h}{\lambda^*}\right)\frac{\partial}{\partial r}\left[\frac{1}{r}\frac{\partial}{\partial r}\left(r\frac{\partial h}{\partial r}\right)\right]\right\} \\ +\frac{\partial}{\partial z}\left(u_z\phi_n\right)=\phi_n g_b-\Psi_d\phi_n, \tag{3.1b}$$

$$\bar{\phi}_n=\frac{1}{h}\int_0^h \phi_n\,\mathrm{d}z, \tag{3.1c}$$

$$u_z=\left(1+\Psi_m\right)g_b\int_0^z \phi_n\,\mathrm{d}\tilde{z} \\ +\frac{\gamma^*}{r}\frac{\partial}{\partial r}\left\{rz\left(\frac{z^2}{6}-\frac{zh}{2}-\frac{h}{\lambda^*}\right)\frac{\partial}{\partial r}\left[\frac{1}{r}\frac{\partial}{\partial r}\left(r\frac{\partial h}{\partial r}\right)\right]\right\}, \tag{3.1d}$$

$$\frac{\partial g_s}{\partial t}=\frac{D}{r}\frac{\partial}{\partial r}\left(r\frac{\partial g_s}{\partial r}\right)-\theta\left(h-h^*\right)\left[DQ_s\left(g_s-g_b\right)\right], \tag{3.1e}$$

$$\mathrm{Pe}h\frac{\partial g_b}{\partial t}=\theta\left(h-h^*\right)\left[\frac{1}{r}\frac{\partial}{\partial r}\left(rh\frac{\partial g_b}{\partial r}\right)+Q_b\left(g_s-g_b\right)-\Upsilon\bar{\phi}_n g_b h \\ -\frac{\mathrm{Pe}\gamma^*}{3r}\frac{\partial}{\partial r}\left\{rg_b\left(h^3+\frac{3h^2}{\lambda^*}\right)\frac{\partial}{\partial r}\left[\frac{1}{r}\frac{\partial}{\partial r}\left(r\frac{\partial h}{\partial r}\right)\right]\right\}\right]. \tag{3.1f}$$

#### 3.1 Transformation of Cell Volume Fraction Term

Before we outline the numerical scheme, we describe the transformation of $\phi_n(r,z,t)$ to $\Phi(r,z,t)$ and subsequently to $\tilde{\Phi}_n(r,\xi,t)=\Phi_n(r,\xi h(r,t),t)$. Taking (3.1b), substituting $z\to\zeta$, then integrating with respect to $\zeta$ from 0 up to $z$ one obtains

$$\frac{\partial \Phi_n}{\partial t}-\frac{\gamma^*}{r}\frac{\partial}{\partial r}\left[\mathcal{H}\int_0^z \phi_n(r,\zeta,t)\left(\frac{\zeta^2}{2}-\zeta h-\frac{h}{\lambda^*}\right)\mathrm{d}\zeta\right]+u_z\phi_n=\Phi_n(g_b-\Psi_d), \tag{3.2}$$

where for convenience we introduce

$$\mathcal{H}(r,t):=r\frac{\partial}{\partial r}\left(\frac{1}{r}\frac{\partial}{\partial r}\left(r\frac{\partial h}{\partial r}\right)\right). \tag{3.3}$$

One can apply integration by parts on the integral within this equation to obtain

$$\int_0^z \phi_n(r,\zeta,t)\left(\frac{\zeta^2}{2}-\zeta h-\frac{h}{\lambda^*}\right)\mathrm{d}\zeta=\Phi_n\left(\frac{z^2}{2}-zh-\frac{h}{\lambda^*}\right) \\ +\int_0^z \Phi_n(r,\zeta,t)(h-\zeta)\,\mathrm{d}\zeta. \tag{3.4}$$

Additionally, observe that $u_z\phi_n$ can be expressed as $u_z(\partial\Phi_n/\partial z)$ and $u_z$ itself can be expressed in terms of $\Phi_n$, to obtain

$$u_z\phi_n = \left((1+\Psi_m)g_b\Phi_n + \frac{\gamma^*}{r}\frac{\partial}{\partial r}\left[\mathcal{H}\left(\frac{z^3}{6} - \frac{z^2 h}{2} - \frac{zh}{\lambda^*}\right)\right]\right)\frac{\partial \Phi_n}{\partial z}. \tag{3.5}$$

Putting the pieces together one obtains

$$\begin{aligned}\frac{\partial\Phi_n}{\partial t} - \frac{\gamma^*}{r}\frac{\partial}{\partial r}\left[\mathcal{H}\left(\Phi_n\left(\frac{z^2}{2} - zh - \frac{h}{\lambda^*}\right) + \int_0^z \Phi_n(r,\zeta,t)(h-\zeta)\,\mathrm{d}\zeta\right)\right]\\ + \left((1+\Psi_m)g_b\Phi_n + \frac{\gamma^*}{r}\frac{\partial}{\partial r}\left[\mathcal{H}\left(\frac{z^3}{6} - \frac{z^2 h}{2} - \frac{zh}{\lambda^*}\right)\right]\right)\frac{\partial\Phi_n}{\partial z}\\ = \Phi_n(g_b - \Psi_d).\end{aligned} \tag{3.6}$$

Now we make the change of variables $(r,z)\to(r,\xi)$ where $z=\xi h(r,t)$. Introducing $\tilde{\Phi}_n(r,\xi,t) := \Phi_n(r,\xi h(r,t),t) = \Phi_n(r,z,t)$, we have

$$\frac{\partial\Phi_n}{\partial z} = \frac{1}{h}\frac{\partial\tilde{\Phi}_n}{\partial\xi}, \tag{3.7a}$$

$$\frac{\partial\Phi_n}{\partial r} = \frac{\partial\tilde{\Phi}_n}{\partial r} - \frac{\xi}{h}\frac{\partial h}{\partial r}\frac{\partial\tilde{\Phi}_n}{\partial\xi}, \tag{3.7b}$$

$$\frac{\partial\Phi_n}{\partial t} = \frac{\partial\tilde{\Phi}_n}{\partial t} - \frac{\xi}{h}\frac{\partial h}{\partial t}\frac{\partial\tilde{\Phi}_n}{\partial\xi}. \tag{3.7c}$$

Similar applies for the partial derivatives of more complex terms in equation (3.6) which have $z$ dependence. For example, applying the transformation to $u_z$ results in

$$\begin{aligned}\tilde{u}_z(r,\xi,t) := u_z(r,\xi h(r,t),t) = (1+\Psi_m)g_b\tilde{\Phi}_n\\ +\frac{\gamma^*}{r}\frac{\partial}{\partial r}\left[\mathcal{H}\left(\frac{\xi^3h^3}{6} - \frac{\xi^2h^3}{2} - \frac{\xi h^2}{\lambda^*}\right)\right] - \frac{\gamma^*}{r}\frac{\partial h}{\partial r}\mathcal{H}\left(\frac{\xi^3h^2}{2} - \xi^2h^2 - \frac{\xi h}{\lambda^*}\right),\end{aligned} \tag{3.8}$$

noting in particular the treatment of the $\partial/\partial r$ term. Upon applying this transformation to the rest of equation (3.6), substituting $\partial h/\partial t$ according to equation (3.1a), and simplifying, one arrives at

$$\begin{aligned}\frac{\partial\tilde{\Phi}_n}{\partial t} - \frac{\gamma^*}{r}\frac{\partial}{\partial r}\left\{\mathcal{H}\left[\tilde{\Phi}_n\left(\frac{\xi^2h^2}{2} - \xi h^2 - \frac{h}{\lambda^*}\right) + h^2\int_0^\xi \tilde{\Phi}_n(1-\bar{\xi})\,\mathrm{d}\bar{\xi}\right]\right\}\\ + \left[(1+\Psi_m)g_b\left(\tilde{\Phi}_n - \xi\tilde{\Phi}_n(r,1,t)\right) + \frac{\gamma^*}{r}\left(\frac{\xi^3}{6} - \frac{\xi^2}{2} + \frac{\xi}{3}\right)\frac{\partial(\mathcal{H}h^3)}{\partial r}\right]\frac{1}{h}\frac{\partial\tilde{\Phi}_n}{\partial\xi}\\ = \tilde{\Phi}_n(g_b - \Psi_d).\end{aligned} \tag{3.9}$$

The integral in the preceding equation is numerically approximated using the trapezoidal rule upon applying a centred finite difference approximation.

It is important to also describe how the initial and boundary conditions for $\phi_n(r, z, t)$ translate to $\tilde{\Phi}_n(r, \xi, t)$. For the initial condition, we obtain

$$\begin{aligned}\tilde{\Phi}_n(r, \xi, 0) &= h(r, 0)\left(\xi^3 - \frac{\xi^4}{2}\right)(1 - 3r^2 + 2r^3)\theta(r - 1) &\quad (3.10a)\\ &= H_0\left(\xi^3 - \frac{\xi^4}{2}\right)(1 - r^2)^4(1 - 3r^2 + 2r^3)\theta(r - 1). &\quad (3.10b)\end{aligned}$$

For the boundary condition,

$$\frac{\partial \phi_n}{\partial r}(0, z, t) = 0, \tag{3.11}$$

upon substituting $z = \zeta$, integrating with respect to $\zeta$ from 0 to $z$, and then swapping the order of integration and differentiation it follows that

$$\frac{\partial \Phi_n}{\partial r}(0, z, t) = 0. \tag{3.12}$$

Expressing this in terms of $\tilde{\Phi}_n$ then results in

$$\begin{aligned}0 &= \frac{\partial \Phi_n}{\partial r}(0, z, t)\\ &= \frac{\partial \tilde{\Phi}_n}{\partial r}(0, \xi, t) - \frac{\xi}{h(0, t)}\frac{\partial h}{\partial r}(0, t)\frac{\partial \tilde{\Phi}_n}{\partial \xi}(0, \xi, t) \qquad (3.13)\\ &= \frac{\partial \tilde{\Phi}_n}{\partial r}(0, \xi, t),\end{aligned}$$

with the last equality due to the boundary condition $\partial h/\partial r = 0$ at $r = 0$. Additionally, given $h = b$ is enforced at $r = R$, then the biofilm is too thin to support any cells, and we can therefore enforce $\phi_n(R, z, t) = 0$ and consequently $\tilde{\Phi}_n(R, \xi, t) = 0$ also. This introduced boundary condition alleviates the need for implementing a one-sided finite difference stencil at this boundary. Observe that for $\xi = 1$ the factor in front of $\partial\tilde{\Phi}_n/\partial\xi$ in the equation (3.9) is exactly zero so that on the biofilm surface the PDE is effectively only evolving radially over the surface.

#### 3.2 Discretisation of the Two-Dimensional Model

The biofilm domain and each of the variables of interest are discretised as follows. Let $r_0 = 0, r_1, \dots, r_{I-1}, r_I = R$ be an equidistant discretisation of the interval $[0, R]$, *i.e.* such that $r_i = i\Delta r$ where $\Delta r = R/I$. Additionally, let $\Delta t$ be some fixed time step size and $t_k := k\Delta t$. Then, let

$$h_i^k := h(r_i, t_k), \quad g_{s,i}^k := g_s(r_i, t_k), \quad g_{b,i}^k := g_b(r_i, t_k). \tag{3.14}$$

Similarly, let $\xi_0 = 0, \xi_1, \ldots, \xi_{J-1}, \xi_J = 1$ be an equidistant discretisation of the interval $[0,1]$, i.e. such that $\xi_j = j\Delta\xi$ where $\Delta\xi = 1/J$. Then, let

$$\Phi_{n,i,j}^{k} = \tilde{\Phi}_n(r_i, \xi_j, t_k) \tag{3.15}$$

(noting the tilde has been dropped from the discretised variable for convenience) and, additionally, let $\phi_{n,i,j}^{k} = \phi_n(r_i, \xi_j h_i^k, t_k)$.

Applying the Crank–Nicolson method to equation (3.1a) governing $h$, using centred finite difference stencils for spatial derivatives, yields the discrete equations

$$\begin{aligned} \frac{h_i^{k+1} - h_i^k}{\Delta t} + \frac{\gamma^*}{6 r_i \Delta r} \Bigg[ & \mathcal{H}_{i+1/2}^{k} \left( (h_{i+1/2}^{k})^3 + \frac{3(h_{i+1/2}^{k})^2}{\lambda^*} \right) - \mathcal{H}_{i-1/2}^{k} \left( (h_{i-1/2}^{k})^3 + \frac{3(h_{i-1/2}^{k})^2}{\lambda^*} \right) \\ & + \mathcal{H}_{i+1/2}^{k+1} \left( (h_{i+1/2}^{k+1})^3 + \frac{3(h_{i+1/2}^{k+1})^2}{\lambda^*} \right) - \mathcal{H}_{i-1/2}^{k+1} \left( (h_{i-1/2}^{k+1})^3 + \frac{3(h_{i-1/2}^{k+1})^2}{\lambda^*} \right) \Bigg] \\ = \frac{1 + \Psi_m}{2} & \left[ \theta(h_i^k - h^*) g_{b,i}^{k} \Phi_{n,i,J}^{k} + \theta(h_i^{k+1} - h^*) g_{b,i}^{k+1} \Phi_{n,i,J}^{k+1} \right], \end{aligned} \tag{3.16}$$

where $r_{i+1/2} := (r_i + r_{i+1})/2 = (i + 1/2)\Delta r$, similarly $h_{i+1/2}^{k} := (h_i^k + h_{i+1}^k)/2$, and lastly

$$\begin{aligned} \mathcal{H}_{i+1/2}^{k} := \frac{r_{i+1/2}}{\Delta r^3} \Bigg[ & \frac{r_{i+3/2}(h_{i+2}^k - h_{i+1}^k) - r_{i+1/2}(h_{i+1}^k - h_i^k)}{r_{i+1}} \\ & - \frac{r_{i+1/2}(h_{i+1}^k - h_i^k) - r_{i-1/2}(h_i^k - h_{i-1}^k)}{r_i} \Bigg]. \end{aligned} \tag{3.17}$$

Observe that taking the half steps with respect to $r$ is essential to ensure that the resulting stencil is only five points wide.

Applying the Crank–Nicolson method to equation (3.9) governing $\tilde{\Phi}_n$, using centred finite difference stencils for spatial derivatives, yields the discrete equations

$$
\begin{aligned}
&\frac{\Phi_{n,i,j}^{k+1}-\Phi_{n,i,j}^{k}}{\Delta t} \\
&\quad -\frac{\gamma^*}{2r^i}\Bigg[\mathcal{H}_{i+1/2}^{k}\left(\Phi_{n,i+1/2,j}^{k}\left(\frac{(\xi_j h_{i+1/2}^{k})^2}{2}-\xi_j(h_{i+1/2}^{k})^2-\frac{h_{i+1/2}^{k}}{\lambda^*}\right)+(h_{i+1/2}^{k})^2\mathcal{I}_{i+1/2,j}^{k}\right) \\
&\qquad\qquad -\mathcal{H}_{i-1/2}^{k}\left(\Phi_{n,i-1/2,j}^{k}\left(\frac{(\xi_j h_{i-1/2}^{k})^2}{2}-\xi_j(h_{i-1/2}^{k})^2-\frac{h_{i-1/2}^{k}}{\lambda^*}\right)+(h_{i-1/2}^{k})^2\mathcal{I}_{i-1/2,j}^{k}\right)\Bigg] \\
&\quad -\frac{\gamma^*}{2r^i}\Bigg[\mathcal{H}_{i+1/2}^{k+1}\left(\Phi_{n,i+1/2,j}^{k+1}\left(\frac{(\xi_j h_{i+1/2}^{k+1})^2}{2}-\xi_j(h_{i+1/2}^{k+1})^2-\frac{h_{i+1/2}^{k+1}}{\lambda^*}\right)+(h_{i+1/2}^{k+1})^2\mathcal{I}_{i+1/2,j}^{k+1}\right) \\
&\qquad\qquad -\mathcal{H}_{i-1/2}^{k+1}\left(\Phi_{n,i-1/2,j}^{k+1}\left(\frac{(\xi_j h_{i-1/2}^{k+1})^2}{2}-\xi_j(h_{i-1/2}^{k+1})^2-\frac{h_{i-1/2}^{k+1}}{\lambda^*}\right)+(h_{i-1/2}^{k+1})^2\mathcal{I}_{i-1/2,j}^{k+1}\right)\Bigg] \\
&\quad +\frac{1}{2}\Bigg[(1+\Psi_m)g_{b,i}^{k}(\Phi_{n,i,j}^{k}-\xi_j\Phi_{n,i,J}^{k}) \\
&\qquad +\frac{\gamma^*}{6r_i}\left((\xi_j)^3-3(\xi_j)^2+2\xi_j\right)\frac{(h_{i+1/2}^{k})^3\mathcal{H}_{i+1/2}^{k}-(h_{i-1/2}^{k})^3\mathcal{H}_{i-1/2}^{k}}{\Delta r}\Bigg]\frac{\Phi_{n,i,j+1}^{k}-\Phi_{n,i,j-1}^{k}}{2h_i^k\Delta\xi} \\
&\quad +\frac{1}{2}\Bigg[(1+\Psi_m)g_{b,i}^{k+1}(\Phi_{n,i,j}^{k+1}-\xi_j\Phi_{n,i,J}^{k+1}) \\
&\qquad +\frac{\gamma^*}{6r_i}\left((\xi_j)^3-3(\xi_j)^2+2\xi_j\right)\frac{(h_{i+1/2}^{k+1})^3\mathcal{H}_{i+1/2}^{k+1}-(h_{i-1/2}^{k+1})^3\mathcal{H}_{i-1/2}^{k+1}}{\Delta r}\Bigg]\frac{\Phi_{n,i,j+1}^{k+1}-\Phi_{n,i,j-1}^{k+1}}{2h_i^{k+1}\Delta\xi} \\
&\quad =\frac{1}{2}\left[\Phi_{n,i,j}^{k}(g_{b,i}^{k}-\Psi_d)+\Phi_{n,i,j}^{k+1}(g_{b,i}^{k+1}-\Psi_d)\right],
\end{aligned}
\tag{3.18}
$$

where $\Phi_{n,i+1/2,j} = (\Phi_{n,i,j} + \Phi_{n,i+1,j})/2$, and similarly $\mathcal{I}_{i+1/2,j}^k = (\mathcal{I}_{i,j}^k + \mathcal{I}_{i+1,j}^k)/2$, where

$$
I_{i,j}^k := \begin{cases} 0 & \text{if } j = 0, \\ \frac{\Delta\xi}{2}\sum_{\ell=0}^{j-1}\left((1-\xi_\ell)\Phi_{n,i,\ell}^k + (1-\xi_{\ell+1})\Phi_{n,i,\ell+1}^k\right) & \text{otherwise.}\end{cases}
\tag{3.19}
$$

Applying the Crank–Nicolson method to equation (3.1e) governing $g_s$, using centred finite difference stencils for spatial derivatives, yields the discrete equations

$$
\begin{aligned}
\frac{g_{s,i}^{k+1}-g_{s,i}^{k}}{\Delta t} &= \frac{D}{2}\Bigg[\frac{r_{i+1/2}(g_{s,i+1}^{k}-g_{s,i}^{k})-r_{i-1/2}(g_{s,i}^{k}-g_{s,i-1}^{k})}{r_i\Delta r^2} \\
&-\theta(h_i^k-h^*)Q_s(g_{s,i}^k-g_{b,i}^k)+\frac{r_{i+1/2}(g_{s,i+1}^{k+1}-g_{s,i}^{k+1})-r_{i-1/2}(g_{s,i}^{k+1}-g_{s,i-1}^{k+1})}{r_i\Delta r^2} \\
&-\theta(h_i^{k+1}-h^*)Q_s(g_{s,i}^{k+1}-g_{b,i}^{k+1})\Bigg].
\end{aligned}
\tag{3.20}
$$

Applying the Crank–Nicolson method to equation (3.1f) governing $g_b$, using centred finite difference stencils for spatial derivatives, yields the discrete equations

$$\begin{aligned}
&\mathrm{Pe}\frac{h_i^k + h_i^{k+1}}{2}\frac{g_{b,i}^{k+1} - g_{b,i}^k}{\Delta t} = \\
&\quad\frac{\theta(h_i^k - h^*)}{2}\Bigg[\frac{r_{i+1/2}h_{i+1/2}^k(g_{b,i+1}^k - g_{b,i}^k) - r_{i-1/2}h_{i-1/2}^k(g_{b,i}^k - g_{b,i-1}^k)}{r_i\Delta r^2} \\
&\qquad + Q_b(g_{s,i}^k - g_{b,i}^k) - \Upsilon\Phi_{n,i,J}^k g_{b,i}^k \\
&\qquad - \frac{\mathrm{Pe}\gamma^*}{3r_i\Delta r}\left(\mathcal{H}_{i+1/2}^k g_{b,i+1/2}^k\left((h_{i+1/2}^k)^3 + \frac{3(h_{i+1/2}^k)^2}{\lambda^*}\right)\right. \\
&\qquad\qquad \left.- \mathcal{H}_{i-1/2}^k g_{b,i-1/2}^k\left((h_{i-1/2}^k)^3 + \frac{3(h_{i-1/2}^k)^2}{\lambda^*}\right)\right)\Bigg] \\
&\quad + \frac{\theta(h_i^{k+1} - h^*)}{2}\Bigg[\frac{r_{i+1/2}h_{i+1/2}^{k+1}(g_{b,i+1}^{k+1} - g_{b,i}^{k+1}) - r_{i-1/2}h_{i-1/2}^{k+1}(g_{b,i}^{k+1} - g_{b,i-1}^{k+1})}{r_i\Delta r^2} \\
&\qquad + Q_b(g_{s,i}^{k+1} - g_{b,i}^{k+1}) - \Upsilon\Phi_{n,i,J}^{k+1} g_{b,i}^{k+1} \\
&\qquad - \frac{\mathrm{Pe}\gamma^*}{3r_i\Delta r}\left(\mathcal{H}_{i+1/2}^{k+1} g_{b,i+1/2}^{k+1}\left((h_{i+1/2}^{k+1})^3 + \frac{3(h_{i+1/2}^{k+1})^2}{\lambda^*}\right)\right. \\
&\qquad\qquad \left.- \mathcal{H}_{i-1/2}^{k+1} g_{b,i-1/2}^{k+1}\left((h_{i-1/2}^{k+1})^3 + \frac{3(h_{i-1/2}^{k+1})^2}{\lambda^*}\right)\right)\Bigg].
\end{aligned} \tag{3.21}$$

Non-Dirichlet boundary conditions for each variable are enforced using one-sided finite difference stencils which are at least second order accurate. For $h$, where there are two boundary conditions at both $r = 0$ and $r = R$, these are enforced in place of stencils at the points $i = 0, 1, I-1, I$. These are enforced implicitly and close the discrete system of equations for the interior points as described above. To avoid the need to implement a biased stencil for the $\tilde{\Phi}_n$ equation at $r = r_1 = \Delta r$ we instead extrapolate these points smoothly from the boundary at $r = 0$ utilising the boundary condition $(\partial\tilde{\Phi}_n/\partial r)(0, \xi, t) = 0$ (this is enforced implicitly in the discrete system of equations).

It would be laborious and uninstructive to explicitly write down the Newton iteration that arises from this system of nonlinear equations. Instead, we refer the reader to our Python class which sets up and solves this system of equations, available at https://github.com/brendanharding/BiofilmLubricationModelSolvers. Once the solution is obtained, the cell concentration $\phi_{n,i,j}^k$ may be recovered from $\Phi_{n,i,j}^k$ via the second order finite difference stencil

$$\phi_{n,i,j}^k = \begin{cases} \theta(h_i^k - h^*)\frac{-3\Phi_{n,i,0}^k + 4\Phi_{n,i,1}^k - \Phi_{n,i,2}^k}{2\Delta\xi h_i^k} & \text{if } j = 0, \\ \theta(h_i^k - h^*)\frac{3\Phi_{n,i,J}^k - 4\Phi_{n,i,J-1}^k + \Phi_{n,i,J-2}^k}{2\Delta\xi h_i^k} & \text{if } j = J, \\ \theta(h_i^k - h^*)\frac{\Phi_{n,i,j+1}^k - \Phi_{n,i,j-1}^k}{2\Delta\xi h_i^k} & \text{otherwise.} \end{cases} \tag{3.22}$$

Additionally, the average cell concentration through each vertical slice is recovered as simply $\bar{\phi}_n(r_i, t_k) = \bar{\phi}^k_{n,i} = \Phi^k_{n,i,J}/h^k_i$.

#### 3.3 Discretisation of the One-Dimensional Model

The discretisation for the one-dimensional model is identical for the equations governing $h$, $g_s$, and $g_b$ with the minor change that $\Phi^k_{n,i,J}$ is replaced by $\Phi^k_{n,i}$ where

$$\Phi^k_{n,i} := \Phi_n(r_i, t_k) = \phi_n(r_i, t_k)h(r_i, t_k) \tag{3.23}$$

in the context of the one-dimensional model. The equation governing $\Phi_n$ is much simpler than in the two-dimensional case, specifically we use (2.1) to obtain the equation

$$\frac{\partial \Phi_n}{\partial t} + \frac{\gamma^*}{3r}\frac{\partial}{\partial r}\left\{\mathcal{H}\Phi_n\left(h^2 + \frac{3h}{\lambda^*}\right)\right\} = \Phi_n(g_b - \Psi_d). \tag{3.24}$$

Applying the Crank–Nicolson method this equation with centred finite difference stencils for spatial derivatives yields the discrete equation

$$\begin{aligned} &\frac{\Phi^{k+1}_{n,i} - \Phi^k_{n,i}}{\Delta t} + \frac{\gamma^*}{6r_i\Delta r}\Bigg[ \\ &\mathcal{H}^k_{i+1/2}\Phi^k_{n,i+1/2}\left((h^k_{i+1/2})^2 + \frac{3h^k_{i+1/2}}{\lambda^*}\right) - \mathcal{H}^k_{i-1/2}\Phi^k_{n,i-1/2}\left((h^k_{i-1/2})^2 + \frac{3h^k_{i-1/2}}{\lambda^*}\right) \\ &+\mathcal{H}^{k+1}_{i+1/2}\Phi^{k+1}_{n,i+1/2}\left((h^{k+1}_{i+1/2})^2 + \frac{3h^{k+1}_{i+1/2}}{\lambda^*}\right) - \mathcal{H}^{k+1}_{i-1/2}\Phi^{k+1}_{n,i-1/2}\left((h^{k+1}_{i-1/2})^2 + \frac{3h^{k+1}_{i-1/2}}{\lambda^*}\right)\Bigg] \\ &= \frac{1}{2}\left(\Phi^k_{n,i}(g^k_{b,i} - \Psi_d) + \Phi^{k+1}_{n,i}(g^{k+1}_{b,i} - \Psi_d)\right). \end{aligned} \tag{3.25}$$

Treatment of boundary conditions and $\Phi^k_{n,1}$ is similar to that described for the two-dimensional model. The resulting discrete system of nonlinear equations may be solved via Newton's method. A Python class for solving this system is available at https://github.com/brendanharding/BiofilmLubricationModelSolvers.
